# Temperate phages limit their own propagation under spatial constraint in biofilms on chitin

**DOI:** 10.1101/2025.11.26.690769

**Authors:** Yixuan Peng, Jacob D. Holt, Triana N. Dalia, Daniel Schultz, Ankur B. Dalia, Carey D. Nadell

## Abstract

Biofilm growth and phage exposure are common to diverse bacterial species. Studying phage-host interaction and population dynamics in biofilms with cellular resolution remains a significant challenge, especially when attempting to recapitulate the natural environments that microbes occupy. Here we study the population dynamics of phage K139 lysogenized and non-lysogenized *Vibrio cholerae* when growing in biofilms on the surface of chitin particles in seawater, replicating key features of *V. cholerae* ecology in the marine environment. We find that lysogenized *V. cholerae*, via spontaneous lytic induction and phage release, kill and displace non-lysogenized bacteria on chitin surfaces. After lysogens become common, however, they can no longer displace remaining non-lysogenized cells. Using a combination of modeling approaches and microscopy experiments, we show that the lysogens’ capacity to displace non-lysogens depends on the ability of phages released by spontaneous induction to reach susceptible non-lysogens. Phage access to non-lysogenized bacterial hosts declines once lysogens become common, and this occurs due to the spatial constraints inherent to biofilm growth as well as to superinfection immunity, which neutralizes phage particles adsorbed to lysogens. Once lysogens comprise the majority of the host population, they can be selected against, because they still incur the cost of spontaneous induction without gaining the benefit of killing non-lysogen cells via phage release. The cost of lytic induction, phage-mediated killing of non-lysogens, and constraints on phage mobility within host bacterial biofilms together yield population dynamics that are consistent with negative frequency dependent selection for lysogenized cells under physiologically realistic growth conditions.

**Significance Statement:** Bacteria often produce and live within biofilm communities in natural environments, where they also encounter many threats including bacteriophages. Here we show how temperate phages can confer a competitive advantage to their hosts via lytic induction and phage release within biofilms of *V. cholerae* on lab-grown marine snow particles. However, the spatial constraints of biofilm architecture and phage-neutralizing superinfection immunity place limits on the extent to which lysogens can competitively displace non-lysogens via phage release.

## Introduction

Temperate phages are defined by their flexibility in using host bacteria as a resource for production of new phage virions (lytic cycle) or as a refuge in which they may propagate their genome vertically within the host bacterium as it grows and divides (lysogenic cycle)(1, 2). A well-studied and expanding list of environmental conditions, including exposure to DNA-damaging agents, high host cell density, and others, can cause a prophage to revert back to the lytic cycle and co-opt the host’s biosynthetic machinery to produce a burst of new phages (3–10). Lytic induction can also occur spontaneously, typically at low rates, within populations of host lysogens that are experiencing conditions that do not cause lytic induction en masse (11–13). The released phages from spontaneous low-frequency induction often do not affect lysogens due to superinfection immunity, but they may contact non-lysogens and either kill them during a lytic infection or otherwise integrate into their genome to produce new lysogens (6, 14, 15). In this manner, temperate phages can displace susceptible cells either by converting them to new lysogens, or by killing them, which not only releases bursts of phage virions but also relieves intraspecific competition for limiting space and growth substrates for other lysogens that have not been induced to the lytic cycle (4, 6, 16–20).

The ability of phage particles to access and exploit host bacteria depends on many factors ranging from receptor compatibility with exposed host surface structures for phage binding, to avoidance of phage defense systems (21–28). On the bacterial community scale, the physical environment can also influence phages’ ability to access hosts (29–33). Barriers to phage transport are particularly relevant in biofilm environments, which many microbes occupy in nature (33–36). Biofilm growth is the process by which diverse microbes immobilize themselves in free-floating groups or in association with surfaces; this mode of growth is thought to be widespread and entails substantial changes in cell physiology, including the defining behavior of secreting an extracellular matrix composed of polysaccharides, proteins, DNA, and potentially other components (37–40). In some cases, phage diffusion can be slowed or halted entirely within biofilms, which leads to altered phage-host population dynamics. Subtle differences in biofilm structure – such as species intermixing and matrix composition – determine whether susceptible bacteria are shielded from phage attacks or sustain phage propagation (41–44).

Understanding how phages propagate among biofilm-dwelling hosts is important for both fundamental and applied perspectives: phages are likely to encounter host bacteria in a biofilm context frequently in natural settings, given the ubiquity of biofilm production across diverse microbial species and environmental conditions (45–47). Additionally, understanding the constraints on phage infection in biofilms is relevant to the growing effort to develop phages as alternative antimicrobials (48–50). While the research community has made significant strides in understanding how biofilm architecture influences phage propagation in recent years (29, 36, 41, 43, 44, 51, 52), some notable hurdles remain. For example, little is known about how well phages released from biofilm-embedded lysogens can propagate through the rest of the host population, particularly in physiologically realistic conditions. Here we study this question using *Vibrio cholerae* growing on chitin particles under flow of artificial seawater, which recapitulates key features of marine snow-associated biofilms in aquatic environments (53, 54).

Using a microfluidic model of chitin particles colonized by fluorescent strains of *V. cholerae*, we tracked the population composition of lysogenized and non-lysogenized bacteria over time. We paired these experiments with well-mixed liquid culture data and simple models of phage-host population dynamics to explore how the marine snow biofilm environment influences phage-bacterial interaction and competition between lysogenized and non-lysogenized hosts. This approach allowed us to determine how biofilm structure influences the propagation of temperate phages released by spontaneous induction, as well as the resulting population biology of lysogenized and non-lysogenized host cells. By addressing these questions, our study explores to what extent phages originating from biofilm-embedded lysogens can spread through host populations and thereby influence competition between bacteria that carry prophages and others that do not. We find that when hosts are confined to chitin surfaces, spatial constraint and superinfection immunity promote coexistence of lysogenized and non-lysogenized bacteria.

## Results

In the marine environments that *V. cholerae* most often occupies, particles of chitinous arthropod exoskeleton are a primary growth substrate (54–58). *V. cholerae* first attaches to and forms biofilms on chitin particles, which leads to upregulation of the machinery required for extracellular digestion via secreted chitinases (54, 55). Our model system is designed to mimic this natural setting and is composed of *Vibrio cholerae* biofilms growing on the surface of chitin particles under active flow of artificial seawater in polydimethylsiloxane (PDMS) flow devices. Experiments were prepared using custom microfluidic devices containing pillars placed in a ‘V’ orientation to trap chitin particles in place (SI Appendix, Fig. S1). The chitin particles were then inoculated with *V. cholerae* under stationary conditions for 30 minutes to allow cells to colonize the chitin surface. After this inoculation, flow was resumed at 0.1 µL/min, corresponding to an average flow velocity of 14 µm/s, which is comparable to the sinking speed of marine snow particles (59–61). All strains of *V. cholerae* were derived from E7946, a pathogenic isolate of the El Tor Ogawa lineage (62). All temperate phage strains were derived from *V. cholerae* bacteriophage K139 from the kappa family of O1 phages (63, 64). As is expected to be the norm among temperate phages with repressor-mediated lysogeny, K139 lysogens exhibit superinfection immunity: phages can adsorb to and inject their genome into lysogens, but this does not lead to lytic infection (64). Lysogens can therefore act as sinks for unproductive adsorption of closely related phages. To distinguish between parental lysogen and parental non-lysogen (i.e., phage susceptible) subpopulations, we used constitutive *mKate22* and *mKO-κ* fluorescent protein constructs integrated in a neutral site in the *V. cholerae* genome. Control experiments indicated that K139 lysogens and non-lysogens have indistinguishable biofilm growth dynamics when they are inoculated independently in monoculture (SI Appendix, Fig. S2); additionally, the *mKate22* and *mKO-κ* fluorescent protein constructs do not influence competition outcomes in co-culture (SI Appendix, Fig. S3). Data were acquired by confocal microscopy of the chitin-attached *V. cholerae* biofilms and analyzed using the image processing framework BiofilmQ (65).

### Population dynamics of lysogens and non-lysogens in marine snow conditions

We aimed first to characterize the competitive dynamics of K139 lysogens and non-lysogenized, phage-susceptible *V. cholerae* cells growing in biofilms on chitin under constant flow, mimicking marine snow conditions in nature. We inoculated chitin particles with varying initial frequencies of lysogens and non-lysogens and then sampled them daily via confocal imaging until the chitin particles were completely consumed by the *V. cholerae* biofilms residing on them. Once chitin particles were fully consumed, the bacteria living on them dispersed, after which population dynamics could no longer be tracked. When lysogens were initially rare, they consistently increased in frequency, suggesting a substantial selective advantage for lysogens against non-lysogens (Fig. 1); lysogens also tended to increase in frequency when they were initiated at intermediate frequencies, although with higher variance in per-day change in lysogen fraction. We speculate that this was due to variation in biofilm size and spatial distribution across replicates. In general, regardless of the initial frequency, as lysogens approached parity in abundance with non-lysogens, their per-day change in frequency became less consistent (Fig. 1A).

**Figure 1:**
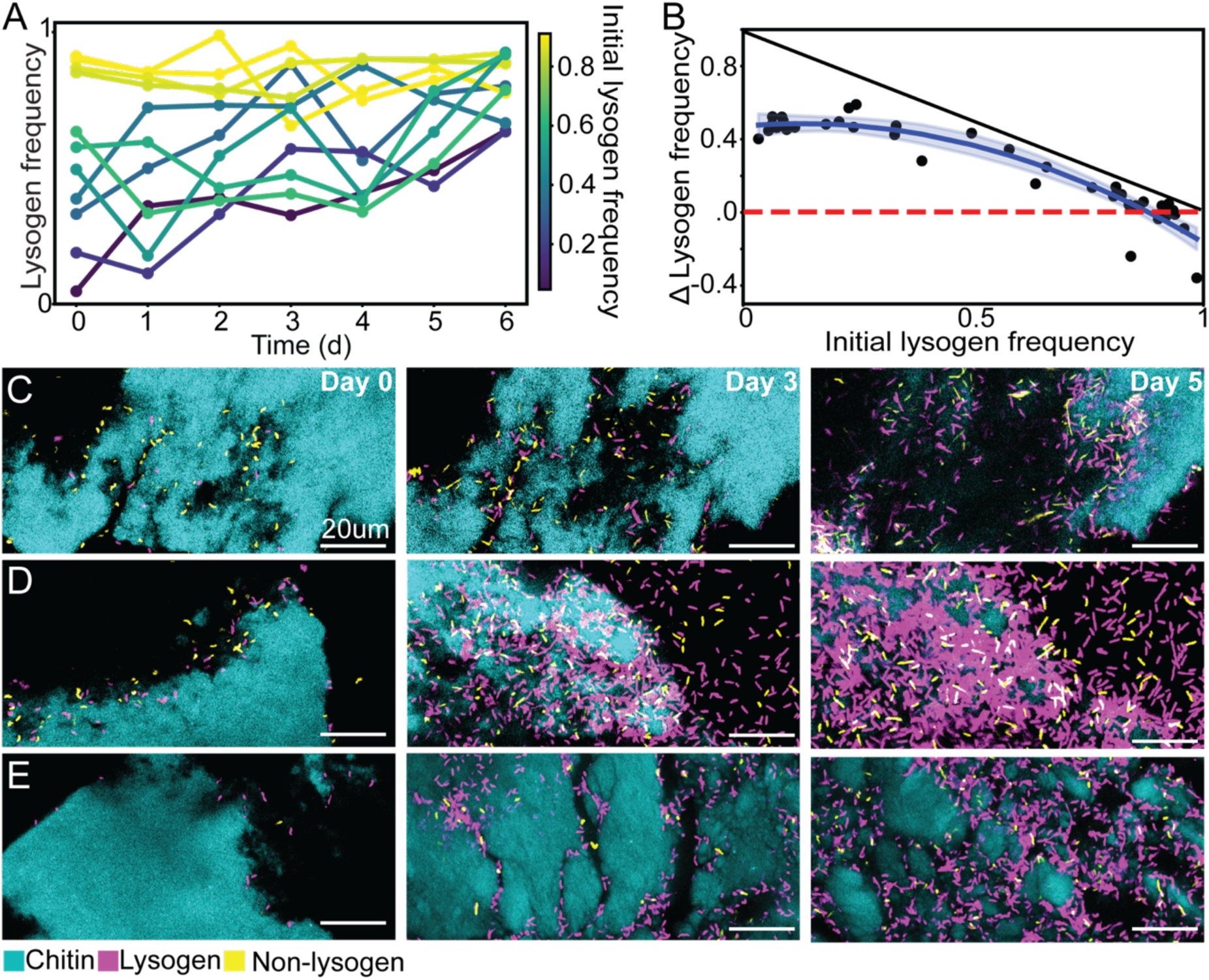
*V. cholerae* K139 lysogens have a fitness advantage in competition with non-lysogens in biofilms on chitin. **(A)** Population dynamics of lysogens and non-lysogens during chitin surface biofilm growth over 6 days (*n* = 11). Lysogens were initiated at a range of initial frequencies, with each trajectory colored according to initial lysogen frequency for visual clarity. **(B)** The relationship between the parental lysogen frequency at day 0 and the change in frequency from day 0 to day 6 (*n* = 39). A quadratic line of best fit is displayed as well (y = 0.47 + 0.22x − 0.89x², R² = 0.89). The dashed red line denotes no change in relative abundance of lysogens over the course of biofilm growth. The black line shows the maximum possible increase in lysogen frequency for each starting condition. **(C, D, E)** Representative 2D images illustrating population dynamics of lysogens and non-lysogens day 0, 3, and 5 at different initial lysogen frequencies. Chitin is displayed in cyan, K139 lysogens in purple, and non-lysogen *V. cholerae* in yellow. The initial lysogen frequencies for panels C, D and E were 0.11, 0.45, and 0.91, respectively.

When lysogens initially comprised the majority of the population, their frequency either did not change, or decreased throughout the experiment. In no instance did we observe lysogens completely displacing non-lysogens in the biofilm populations; overall the population dynamics across experimental runs implied the operation of negative frequency-dependent selection for lysogens in competition with susceptible cells, with lysogens approaching an equilibrium frequency of ∼0.88 (95% confidence interval: [0.85, 0.91]) (Fig. 1B) (66, 67). These results in turn suggested that phage release by spontaneous induction can benefit the remaining lysogen population by removing non-lysogens from the chitin surface, but that this benefit might depend on the frequency of lysogens in the system (16). Importantly, this trend was specific to the biofilm environment; in well-mixed liquid culture controls, lysogens were positively selected regardless of initial frequency (SI Appendix, Fig. S4A). This comparison implies an important role of spatial structure and biofilm architecture underpinning the population dynamics observed in the chitin experiments. Another potential explanation is that phages cannot infect some susceptible cells due to a change in host physiology in the biofilm context, though additional experimental argues against this possibility in showing that widespread death of the susceptible cell population is contingent on infectious phage release from lysogens (see Fig. 3 below).

Control experiments indicated that the frequency of newly-produced lysogens in our experiments was 0.015 (range: [0, 0.02]), and the frequency of *de novo* K139-resistant *V. cholerae* mutants was also 0.015 (range: [0, 0.04]) (SI Appendix, Fig. S5). It is important to note that the ΔK139 mutation used to remove the prophage in our non-lysogen strain also removes the attP site to which K139 integrates when lysogenizing a host. This very likely reduces the potential for phage-susceptible cells in our experiments to become lysogenized after K139 infection, but it also increases the clarity of the distinction between the lysogens and non-lysogens (63, 64). Altogether, these control experiments illustrate that most phage infections were lytic, and that the expansion of the initial lysogen sub-population was mainly due to outgrowth of previously existing lysogens rather than novel lysogenization of phage-susceptible cells. Furthermore, selection for *de novo* phage resistance was not strong on the time scale of these experiments and could not by itself explain why cells with the non-lysogen fluorescent marker remained present through the full duration of the experiments.

Our findings suggest that lysogens gain a frequency-dependent advantage through local phage release that selectively removes nearby susceptible competitors, but only up to a limit determined by spatial constraints in the system and the relative abundance of lysogens, which adsorb and neutralize phages. When lysogens become abundant, reduced opportunities for infection limit phage spread, causing their relative fitness to decline, a pattern consistent with selection against lysogens when they are at high frequency. To test this interpretation, in the following sections we use a simple model and additional experiments to study the mechanistic underpinning of the results presented thus far.

### Lysogens block phage access to susceptible cells in a simple percolation model

Our experiments above document that *V. cholerae* K139 lysogens, when rare or moderate in relative abundance, have a competitive advantage over phage-susceptible non-lysogens in biofilms on chitin. At high frequency, however, lysogens have no advantage and often decrease in frequency relative to non-lysogens. The benefit of partial lytic induction within the lysogen population (removal of competing non-lysogens) must exceed the cost of induction (death of a fraction of the lysogen population) in order for lysogens to obtain an overall fitness advantage in competition with non-lysogenized, susceptible cells. The experimental results in Figure 1 indicate that the balance of this cost and benefit for lysogens changes as they become more abundant. In particular, superinfection immunity in K139 lysogens leads to non-productive adsorption by K139 virions (64) (SI Appendix, Fig. S6). In combination with spatial constraints within biofilm environments and loss of released phage to the flow environment, phage virion adsorption to lysogens might reduce phage mobility to the extent that virion particles cannot reach susceptible cells that are distributed at low abundance in the population. Here we use a simple percolation model to assess the validity of this inference and to study the limitation on susceptible host access by phages as a function of increasing lysogen frequency.

The model implements cells that are fixed in place on a lattice and designated as uninfected non-lysogens, infected non-lysogens, or lysogens (*i.e.*, phage-adsorbing but immune). As we observed limited or no clustering of lysogenized and susceptible *V. cholerae* in our experiments (SI Appendix, Fig. S7), each simulation run was initiated with a random distribution of lysogen and non-lysogen cells, and infection was assumed to begin from a single location in the center of the simulation space. The model tracks the fraction of susceptible cells that are within continuous contact with each other from the initial site of phage infection, with lysogens acting as barriers to infection propagation. To isolate the effect of lysogen frequency on phage access to susceptible hosts, phage replication and diffusion, as well as bacterial cell growth, division, and movement, were not implemented. The two-dimensional version of the model in Figure 2 illustrates the principle that increasing lysogen frequency restricts infection spread by limiting the number of susceptible cells in continuous contact with each other. For the full simulation runs, we extended the model environment to a three-dimensional lattice, which better reflects the observed depth of chitin-bound biofilms seen in our experiments. We calculated the proportion of infected susceptible cells as a function of initial lysogen frequency across 100 replicate simulations for each initial condition. Figure 2B illustrates that the fraction of infected susceptible cells declines precipitously when lysogens comprise 85-90% of the bacterial population.

**Figure 2.**
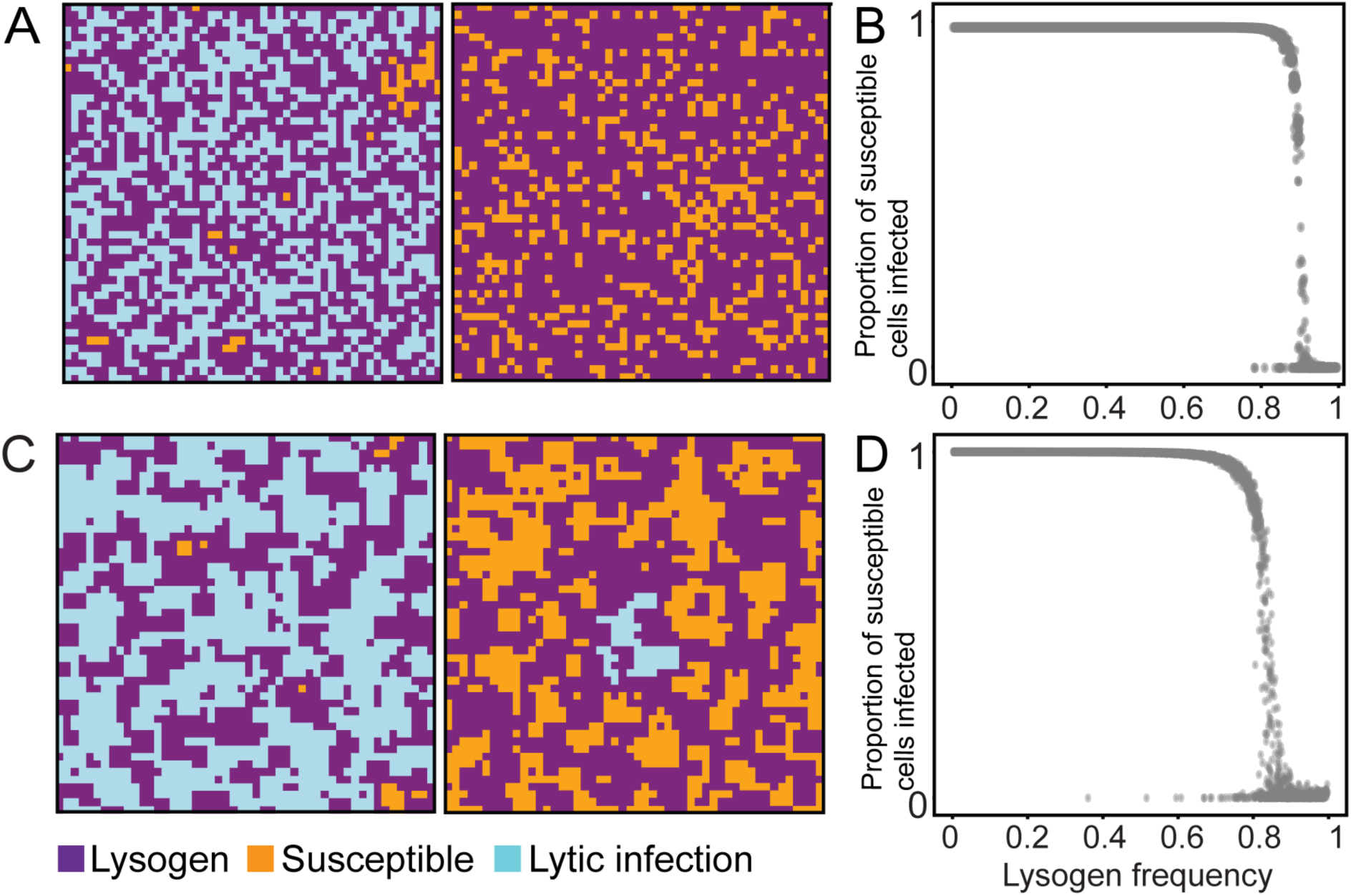
Constraints on infection propagation in a simple percolation model. (A) An illustration of infection spread (left) or containment (right) in a two-dimensional space with randomly distributed lysogens and susceptible cells. (B) Proportion of infected susceptible cells as a function of lysogen frequency for simulations with randomly distributed lysogens and susceptible cells. (C) An illustration of infection spread (left) or containment (right) in a two-dimensional space with clustered distributions of lysogens and phage-susceptible cells. (D) Proportion of infected susceptible cells as a function of lysogen frequency for simulations with clustered distributions of lysogens and susceptible cells.

To account for cases in which lysogens and non-lysogens might have been clustered by clonal outgrowth, we ran additional simulations that were initialized with patchy distributions of the two cell types (Fig. 2C); we also explored the impact of open space between groups of lysogens and susceptible cells, in which phage infection could or could not jump from one patch of cells to another (SI Appendix, Fig. S8). The results from these runs yielded a similar relationship between lysogen frequency and the fraction of infected susceptible cells as observed above, noting that susceptible cell infection declined at somewhat lower lysogen frequencies relative to simulations with random initial distributions of lysogens and susceptible cells. Some simulation runs initiated with clusters of lysogens and non-lysogens also exhibited drops to zero susceptible infection at lower initial lysogen frequencies, which was due to local blocking in which clusters of lysogens trapped infection propagation near the point of origin (Fig. 2D). These results are analogous to well-established theory of herd immunity in the context of infectious disease transmission, and they are consistent with the idea that phages released from spontaneously induced lysogens no longer efficiently infect susceptible cells after lysogens comprise a sufficiently large fraction of the host population (68–71).

In addition to the heuristic percolation model, we also implemented a reaction-diffusion model to study the population dynamics of lysogens, non-lysogens, and free phages, parameterized experimentally where feasible. This model could readily recapitulate the experimental finding that lysogens and non-lysogens tend toward coexistence under spatial constraint, with selection favoring lysogens when they are initially rare or moderate in relative abundance, and selection against lysogens when they are initially common (SI Appendix, Methods and Fig. S9A,B). A sensitivity analysis indicated that the selection regime is contingent on the adsorption rate of phages to lysogens and non-lysogens, and to a lesser degree on lysogens’ lytic induction rate (SI Appendix, Fig. S9C).

Based on the experimental observation that lysogens decrease in frequency when initially in the majority of the population, we infer that the cost of spontaneous induction to produce phages – i.e., cell death – is high enough that when phages are no longer eliminating susceptible cells from the population, lysogens incur a net fitness disadvantage in competition with non-lysogens. In the following section we assess this inference experimentally.

### Lytic induction cost imposes a fitness defect in the absence of phage-mediated killing

Our results thus far suggest that spontaneous lytic induction among K139 lysogens, leading to cell death and phage release, confers a competitive benefit to remaining lysogens by killing off susceptible cells. Once lysogens reach a sufficiently high fraction of the population, the cost of lytic induction is no longer outweighed by the benefit of non-lysogen killing, because few released phages can reach susceptible cells in the midst of a large number of phage-adsorbing lysogens. This leads to a net fitness disadvantage among lysogenized cells that yields mixed populations of lysogens and non-lysogens at steady state. We next sought to test the assumption that phage release is responsible for non-lysogen cell death, as well as the assumption that the cost of prophage induction can be sufficient to explain selection against lysogens when they are common.

To confirm that the competitive advantage of lysogens at low frequencies was mediated by phage-induced lysis of susceptible cells, we measured the degree of cell death in the phage-susceptible strain in the presence of lysogens that produce functional K139 phages after lytic induction. To visualize cell death, we used SYTOX Green, a membrane-impermeable stain that fluoresces upon binding to DNA. Because SYTOX Green only diffuses through compromised membranes, it selectively labels dead or damaged cells. In co-culture experiments including susceptible cells and lysogens harboring K139 prophages, imaging of SYTOX-stained biofilms indicated pervasive phage-susceptible cell death when lysogens were initiated at low frequency (Fig. 3A-B).

**Figure 3:**
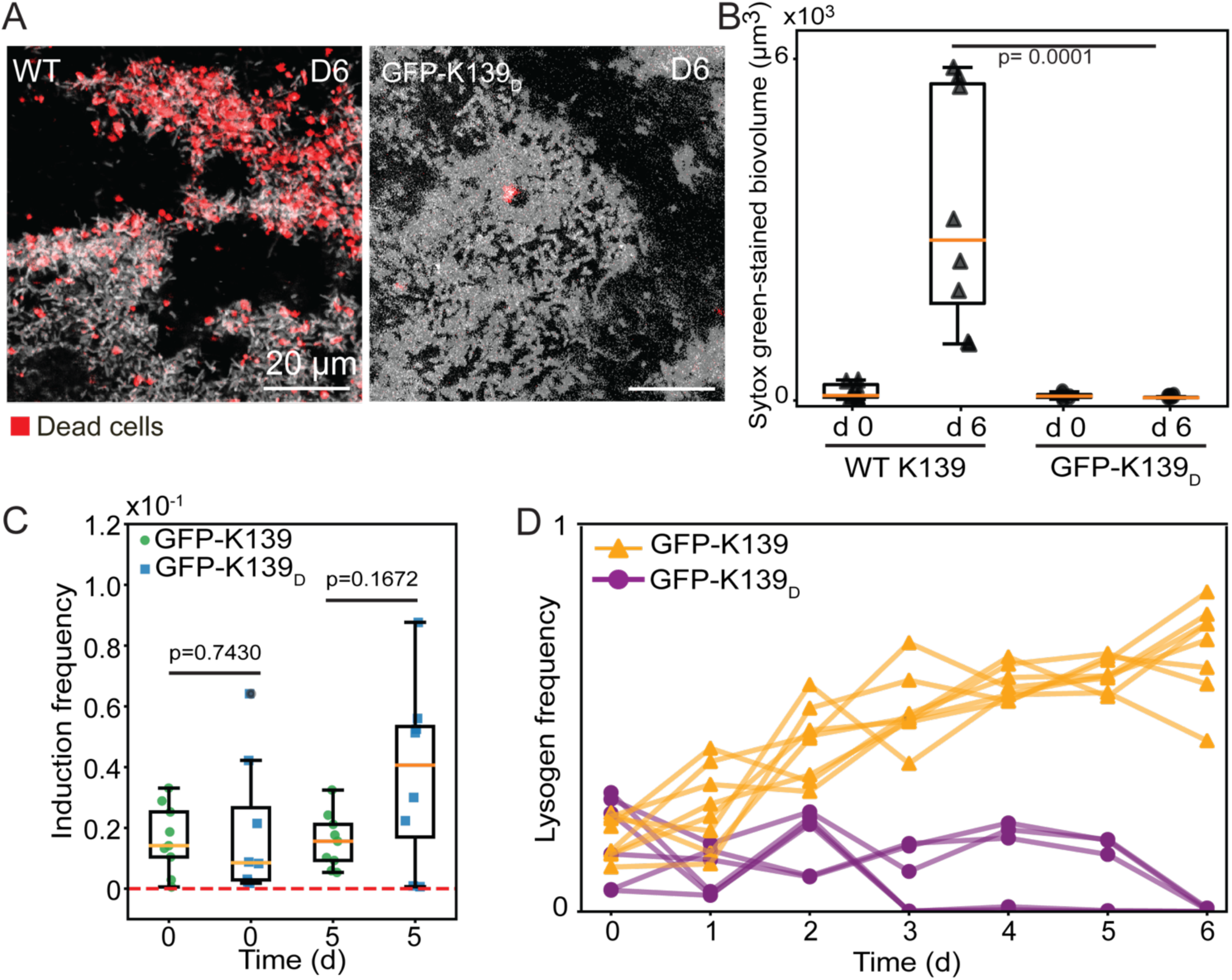
Lytic induction of K139 lysogens incurs a substantial cost to *V. cholerae* lysogens due to cell lysis, and this cost is only offset when phage particles can cause new infections. (**A)** Representative 2D images of *V. cholerae* biofilms on chitin from day 6 co-cultures of susceptible *V. cholerae* with lysogens carrying either K139 (left) or the defective phage variant GFP-K139_D_ (right). For clarity here, the chitin is not shown; live bacteria are shown in grey, while dead cells are stained with SYTOX and shown here in red. **(B)** Quantification of killed susceptible hosts on day 0 and day 6 after co-culture with lysogens carrying WT K139 or GFP-K139_D_ (*n* = 8). **(C)**. Boxplots showing the observed frequency of lytic induction at day 0 and day 5 for lysogens carrying either GFP-K139 (*n* = 8) or GFP-K139_D_ (*n* = 8). **(D)** Time-course of lysogen frequency changes over 6 days, comparing lysogens with functional (orange; *n* = 7) and GFP-K139_D_ (purple; *n* = 6) phages in competition with susceptible population. Lines and dots represent data from each chamber, with replicates each obtained by averaging technical replicates from 2-3 chitin particles. p-values noted in panels B and C were derived from Mann–Whitney U tests.

Because cell death can occur for a variety of reasons during biofilm growth, we sought to isolate the killing effect of phages released from lytic induction by lysogenized *V. cholerae* (72–74). To parse the importance of phage activity in the experiment described above, we constructed two K139 derivative strains with capsid GFP translational fusions that undergo spontaneous lytic induction from host lysogens at the same rate as the parental K139 (Fig. 3C). GFP-K139 produces phage virions that retain their infectivity and was constructed by generating a strain that carries an orf18-GFP translational fusion expressed via its native promoter at an ectopic locus. This configuration allows incorporation of a subset of GFP-tagged proteins into otherwise wild-type K139 capsids, which produces functional and fluorescently labeled phages. The other K139 derivative, denoted GFP-K139_D_ (infection defective), carries a GFP fusion to Orf18 at its native locus within the prophage genome (with no ectopic construct), and it produces GFP-fluorescent virions that are unable to cause new lytic infections in susceptible cells. We inferred that phages released by the GFP-K139_D_ strain were non-infectious because the GFP-labeled version of the Orf18 capsid protein prevents phage adsorption to the host surface when not accompanied by a wild type Orf18 allele in the same strain background (63, 75, 76). We confirmed that induction in GFP-K139_D_ lysogens does lead to lytic release of virions by quantifying phage fluorescent signal through imaging, but the phages released by GFP-K139_D_ do not form plaques on phage-susceptible *V. cholerae* (Fig. 3C, SI Appendix, Fig. S10). When we co-inoculated susceptible *V. cholerae* with lysogens carrying GFP-K139_D_, we observed little to no death of susceptible cells at day 6 (Fig. 3A-B), confirming the role of viable lytic phage infections in killing phage-susceptible cells.

The experiments above directly confirm that lysogens eliminate a substantial fraction of the non-lysogens in the system via release of phages following spontaneous lytic induction. The infection-defective GFP-K139_D_ phage variant also allowed us to assess the outcome of competition between lysogens and susceptible non-lysogens when phage-mediated killing of susceptible cells is toggled on (lysogens carry GFP-K139) or off (lysogens carry GFP-K139_D_). Both the GFP-K139 and GFP-K139_D_ lysogen strains spontaneously induce to lytic phage production at comparable rates (Fig. 3C), such that the cost of lytic induction is held constant, but only phages produced by GFP-K139 confer a potential benefit to lysogens by killing off phage-susceptible cells.

By monitoring the competitive dynamics of susceptible non-lysogens in co-culture with lysogens that release infectious phages, versus lysogens that release non-infectious phages, we could assess the assumption that the cost of spontaneous lytic induction yields a net fitness defect to lysogens when the phages they release cannot kill susceptible hosts. To make this comparison we repeated the competition experiments from Figure 1, focusing on the case in which lysogens are initiated at low frequency, when the benefit of phage release and susceptible cell killing is most pronounced. As observed for lysogens carrying WT K139 prophages Figure 1, GFP-K139 lysogens consistently increase in frequency in competition with susceptible non-lysogens (Fig. 3D). By contrast, GFP-K139_D_ lysogens never increase in frequency despite ordinarily favorable starting conditions (i.e., lysogens in the minority of the population), and by the end of these experiments on day 6, GFP-K139_D_ lysogens were eliminated from the population altogether in all runs of the experiment (Fig. 3D; SI Appendix, Fig. S11). Similarly, in shaken liquid culture experiments, GFP-K139_D_ lysogens were negatively selected from any initial frequency in competition with susceptible non-lysogens (SI Appendix, Fig. S4B). This outcome confirms that the cost of spontaneous lytic induction is sufficiently high to produce a net fitness disadvantage among lysogens when the phages they release have no effect on susceptible host bacteria.

Collectively, our results establish that lysogens can partially displace phage-susceptible strains via spontaneous lytic induction and release of phages that go on to kill off non-lysogenized cells in the population. However, lysogens do not fix in the population; that is, they are negatively selected when they are at high relative abundance. At low lysogen frequencies, the ability to eliminate phage-susceptible competitors via phage-induced lysis outweighs the population cost of prophage induction. As lysogens become more prevalent, phage transmission is limited by spatial constraints inherent to biofilm growth and superinfection immunity of abundant lysogens; but the costs associated with prophage induction are still incurred. As a result, lysogens are negatively selected when they are common.

## Discussion

Using co-culture experiments including K139-lysogenized and non-lysogenized *Vibrio cholerae* on chitin, we demonstrated that spontaneous lytic induction of the K139 prophage leads to substantial killing of phage-susceptible cells, which in turn allows remaining lysogens to occupy more space on the chitin surface. A simple percolation model supports the idea that once lysogens occupy the majority of the population, phage virions released from spontaneously inducing lysogens no longer efficiently access non-lysogenized, phage-susceptible hosts. Because K139 prophages undergo lytic induction at a relatively high basal rate, they impose a significant cost on the lysogen population; when released phages no longer reach and kill susceptible cells, this cost of induction leads to a net disadvantage for lysogens and prevents them from reaching fixation. In analogous experiments in liquid culture, where phage-host cell interaction is homogenized, lysogens increase in frequency from any starting condition. This contrast highlights the importance of realistic growth conditions: due to the combination of susceptible cell death by lytic phage infection, the cost of spontaneous lytic induction by prophages, and spatial constraints on phages’ access to susceptible host cells when lysogens are abundant, lysogens tend to coexist with non-lysogens rather than displacing them.

Our results offer a direct illustration of the limits of phage spread in mixed populations of lysogens and non-lysogens when phages are neutralized by superinfection immunity; they also reinforce the adaptive rationale for superinfection exclusion mechanisms that some temperate phages have evolved to avoid neutralization on lysogens’ surface. Superinfection exclusion - distinct from superinfection immunity – refers to mechanisms that some temperate phages have evolved to prevent attachment of released virions to the surface of other lysogens. Taylor et al. (2025) recently discovered that JBD26 prophages achieve superinfection exclusion by producing the pilZ-interacting protein Zip. Zip leads to reduction in type IV pilus length and total count, preventing phage adsorption to their pilus binding sites. This mechanism, by preventing JBD26 phage progeny from re-adsorbing to lysogens, avoids self-neutralization and maintains a large pool of infectious virions for horizontal spread (77). Our results in this paper offer a direct illustration, in physiologically realistic conditions, of the fitness defect incurred in the absence of mechanisms that prevent phage adsorption to previously lysogenized host bacteria.

The observations in this study are consistent with the concept of herd immunity that has been extensively studied in the infectious disease ecology literature (78–80). Prior work has similarly found that sufficiently high fractions of host bacteria housing CRISPR-based resistance to phage T7 can prevent the phage from accessing the full host population (81). In the system explored here, lysogenization of host bacteria by the temperate phage itself – as opposed to a dedicated, host-evolved phage defense system – imposes limits on its own propagation. Our results also corroborate prior theoretical work in the biofilm literature, which has found that refugia in which some bacterial cells remain unexposed to phages are crucial to coexistence of phages and susceptible hosts (36, 41, 82). These refugia can emerge spontaneously due to biofilm architecture that limits phage diffusion, but in the instance described here, refuge from phage exposure emerges when lysogens become common enough to prevent access to susceptible hosts by phages released by spontaneous induction.

Finally, our results speak to an important reason why bacteria with and without prophages should tend to coexist with one another. The features of our system that lead to coexistence of lysogens and non-lysogens – basal spontaneous induction of lysogenized hosts, phage-neutralizing superinfection immunity by lysogens, and spatial constraints on phage diffusion in biofilms – are common to many temperate phages and biofilm environments (12, 36, 43, 83). Natural variation is expected, though: one recent study reported two novel *Pseudomonas aeruginosa* phages that specifically target the extracellular matrix polysaccharide Psl to infect hosts (84, 85). For any examples in which temperate phages have evolved mechanisms to relieve constraints on their diffusion in biofilms, we might expect them to more effectively lysogenize the entire host population (43, 86, 87). Another important consideration is that our study does not address the fact that many natural biofilm systems are thought to contain dozens or hundreds of different species (88–90). The presence of numerous other species that are not susceptible to a given phage, for example, could impose additional barriers to phage diffusion that would limit the ability of spontaneously induced prophages to alter the community composition (44, 91, 92). Further increases to ecological realism in the design of future experiments will be important to assess the generality of the results reported here.

## Methods Summary

All bacterial strains were derived from *V. cholerae* E7946, and all temperate phage strains were derived from *V. cholerae* bacteriophage K139. Biofilm cultivation was performed using PDMS microfluidic devices, and imaging was performed using a Zeiss 880 point-scanning confocal microscope with a 40x 1.2NA water objective. Data processing and analysis were performed using the BiofilmQ framework in MatLab. Python was used for statistical analysis and figure panel generation. Detailed methods are available in the SI Appendix.

## Data, Materials, and Software Availability

All study data are included in the article and the SI Appendix.

## Acknowledgments

The authors extend their thanks to Sofia Garcia, Tapan Goel, Jacopo Marchi, iela Santiago, Joshua Weitz, Jay Winans, and the Dartmouth microbiology community for feedback on this project. This work was supported by the Simons Foundation (Award 826672 to CDN), the National Institute of General Medical Sciences (grant R35GM151158 to CDN, and grant R35GM128674 to ABD), NSF grants PHY-2412766 and DMS-2527337, and the U.S. Department of Energy grant DE-SC0026232.

## Author Contributions

CDN conceived, funded, and supervised the study. YP, JDH, DS, ABD, and CDN designed models and experiments. YP performed experiments and modeling, analyzed data, and generated the figures. TND and ABD provided key reagents and strains. YP and CDN wrote the paper. All authors contributed to manuscript editing.

## SI Appendix

### Materials and Methods

#### Bacterial strains and growth

The *Vibrio cholerae* strains used in this study were derived from strain E7946. Fluorescent protein expression constructs under IPTG-inducible expression were generated in this study and in prior work using standard molecular biology techniques, including by allelic exchange or by natural transformation on chitin as previously described (93). For strains generated by allelic exchange, Kanamycin 100 μg/mL and Polymyxin 50 μg/mL were used to select for *V. cholerae* after mating with *E. coli*, and *sacB* counter-selection on plates containing 10% sucrose was then used to select for cells that excised the plasmid. Correct strains were confirmed by PCR and sequencing. Cultures were grown overnight in lysogeny broth (LB) supplemented with 100 µM isopropyl β-D-1-thiogalactopyranoside (IPTG) to induce fluorescent protein expression. Colony forming units (CFU) were measured using plain LB plates. Plaque assays were performed using plates prepared with 10 g/L Tryptone, 8 g/L NaCl, 1 g/L yeast extract, 10 g/L agar, 1 g/L glucose, and 2 mM CaCl_2_. Double-layer soft agar was prepared with 10 g/L Tryptone, 8 g/L NaCl, 1 g/L yeast extract, 7 g/L agar, 1 g/L glucose, and 2 mM CaCl_2_ to facilitate phage adsorption. See SI Appendix, Table S1 for a full list of strains and reagents used in this study.

#### Microfluidic device assembly

Microfluidic chambers for biofilm culture were made using polydimethylsiloxane (PDMS) and standard soft lithography techniques. After mixing the PDMS base and curing agent at a 10:1 ratio, the mixture was poured into a silicon wafer mold and cured at 65°C overnight. Following curing, inlet and outlet ports were added to the PDMS blocks using a biopsy punch press. The PDMS chambers were then bonded to #1.5 glass coverslips (36 mm x 60 mm) using room air plasma treatment. Continuous flow within the microfluidic devices was maintained using Harvard Apparatus Pico Plus syringe pumps mounted with 1 mL Brandzig plastic syringes. The syringes were fitted with 27-gauge needles and connected to #30 Cole Parmer PTFE tubing (inner diameter: 0.3 mm). The inlet tubing was attached to the syringe, while the outlet tubing was directed into a petri dish for waste collection.

#### Fluorescence microscopy

Fluorescence imaging was performed using a Zeiss 880 point-scanning confocal microscope with a 40x/1.2 N.A. water objective. GFP-labeled phages and SYTOX Green (dead cell stain) were excited using a 488-nm laser line, mKO-κ was excited using a 543-nm laser line, and mKate22 was excited using a 594-nm laser line. Chitin flakes, which exhibit blue autofluorescence, were excited using a 405-nm laser line. The microscope hardware was controlled using ZEN Black software (Carl Zeiss) for the LSM 880 system.

#### Biofilm growth conditions

For microfluidic experiments, cultures were grown in artificial seawater (ASW) supplemented with 100 µM IPTG and chitin flakes derived from shrimp shells (Sigma). Prior to introduction into microfluidic devices, chitin flakes were washed twice with 70% ethanol, followed by five washes with phosphate-buffered saline (PBS), and a final wash with artificial seawater (ASW) to ensure removal of residual contaminants. For liquid culture experiments, artificial seawater was used as the base medium, supplemented with 0.5% N-acetylglucosamine (GlcNAc). The defined artificial seawater (DASW) used in this study consisted of the following components: 234 mM NaCl, 27.5 mM MgSO_4_, 1.5 mM NaHCO_3_, 4.95 mM CaCl_2_, 5.15 mM KCl, 0.07 mM Na_2_B_4_O_7_, 0.05 mM SrCl_2_, 0.015 mM NaBr, 0.001 mM NaI, 0.013 mM LiCl, 0.187 mM K2HPO4, and 0.1M HEPES solution.

For all chitin-related experiments, washed chitin flakes were suspended in DASW prior to loading and introduced into the microfluidic devices using a pipette. Cultures of each bacterial strain were resuspended in DASW and adjusted to an optical density (OD_600_) of 0.4. For monoculture experiments, 10 µL of a single strain was inoculated into each well of the microfluidic device using a micropipette. For competition experiments, cultures of different strains were mixed at specified ratios immediately prior to inoculation, and 10 µL of the mixed suspensions were introduced into the microfluidic chambers. Following inoculation, the microfluidic devices were left undisturbed for 1 hour at room temperature (22°C) to allow for bacterial colonization of chitin surfaces. The starting populations for each replicate of the control and competition experiments contained 6,000-8,000 cells in total. After this initial incubation period, artificial seawater was introduced to the chambers continuously at a rate of 0.1 µL/min. All experiments were conducted at 22°C.

#### Phage purification and quantification

To produce phages for use in the experiments, a lysogen culture was grown overnight, centrifuged, and filtered through a 0.22 µm membrane to remove bacterial cells. The resulting filtrate was plated on a lawn of susceptible *V. cholerae* using the traditional double layer soft agar plaque assay with strain CNV 272 as the host (94). A single plaque was isolated, resuspended, and used to infect a fresh culture of susceptible cells. After another overnight incubation, the culture was centrifuged and filtered through a 0.22 µm membrane to obtain phage lysate. For phage titer quantification using plaque assays, collected samples were treated with chloroform, vortexed, and allowed to sit for 5 minutes. A 100 µL aliquot of the supernatant was mixed with 100 µL of *V. cholerae* overnights culture in molten soft agar and poured onto LB agar plates supplemented with 10 mM MgSO_4_ and 1 mM CaCl2 to support phage infection. After overnight incubation at 37°C, phage plaques were enumerated to calculate phage titer.

#### Determining the frequency of newly formed lysogens and *de novo* phage-resistant mutants

Following six days of biofilm growth in microfluidic chambers containing *Vibrio cholerae* lysogens (*mKate22*-labeled) and susceptible non-lysogens (*mKO-κ*-labeled), biofilms were washed out by flushing each chamber with 200 µL sterile phosphate-buffered saline (PBS). The resulting cell suspension was serially diluted and plated on LB agar containing IPTG to obtain well-isolated colonies. Colonies expressing the susceptible non-lysogen fluorescent marker (mKO-κ⁺) were identified using fluorescence microscopy and randomly selected for further analysis (≥50 colonies per condition). Colonies were picked and inoculated into LB medium in 96-well plates and grown overnight. Overnight cultures were then used as host lawns for plaque assays with wild-type K139 phage. Colonies that supported plaque formation were classified as susceptible. Colonies that did not support plaque formation could represent either resistant mutants or newly formed lysogens and were subjected to additional testing. These isolates were regrown overnight, and supernatants were collected following chloroform treatment and spotted onto lawns of a K139-susceptible host strain (CNV272) to assess phage production. Colonies whose supernatants produced plaques were classified as newly formed lysogens, whereas colonies that neither supported plaque formation nor produced phages were classified as resistant mutants. The frequencies of susceptible cells, newly formed lysogens, and resistant mutants were calculated as the number of colonies in each category normalized to the total number of colonies screened.

#### Liquid competition experiments

Overnight cultures of K139 lysogens and susceptible non-lysogen cells were diluted in artificial seawater supplemented with 0.5% GlcNAc to OD_600_ = 0.4. The cultures were then mixed at varying ratios and incubated at room temperature with shaking at 250 rpm. To calculate population composition, aliquots were imaged at two time points: immediately after mixing (0 hours) and after 40 hours of incubation. At each time point, 10 µL of the culture was pipetted onto a microscope slide and covered with an agar pad to immobilize the cells for microscopy. End-point strain frequencies were determined using the constitutive fluorescent protein constructs that distinguished each strain.

#### Assessing superinfection immunity

To assess the adsorption kinetics of K139 phages to lysogenized *V. cholerae* populations under well-mixed conditions, we performed time-course assays measuring free phage titers. Cultures of K139-lysogenized *V. cholerae* were grown in artificial seawater supplemented with 0.5% GlcNAc at 37 °C with shaking until mid-exponential phase. K139 phages were then added to the *V. cholerae* culture at an initial concentration of 1×10^5^ PFU/mL. In a control experiment, phages were added at the same concentration to otherwise identical media without bacteria present. The phage-host co-cultures and phage-only control cultures were incubated at 37°C with shaking at 250 rpm in 10 mL tubes. Samples were collected at 0, 30, 60, 120, 180, and 300 minutes, and the concentration of free phage particles (PFU/mL) was determined by traditional double layer plaque assay.

#### Dead cell staining with SYTOX Green

To assess cell viability, SYTOX Green nucleic acid stain was used to label dead cells. A stock solution of SYTOX Green was diluted in artificial seawater to a final concentration of 5 µM. The artificial seawater and SYTOX green staining solution was continuously perfused into the microfluidic device at a flow rate of 0.1 µL/min.

#### One step growth curve experiment

Burst size was determined after carrying out a one-step growth experiment performed as previously described (95, 96) (SI Appendix, Fig. S12). Briefly, phage lysate was diluted and added to a diluted bacterial overnight culture to obtain a low multiplicity of infection (MOI, ratio of phages to hosts). After a brief adsorption step for 10 min, aliquots were sampled at 5 min intervals for 1 h, and phage titers were quantified by traditional double layer plaque assays. Plotting phage titer over time using these initial conditions tracks the rise in phage titer to a plateau following the release of phages from one round of host infection, providing estimates for the latent period between host adsorption and phage burst, as well as the number of phages per burst event.

#### Lysogen Induction rate assay

To quantify the proportion of lysogenized *V. cholerae* cells undergoing spontaneous phage induction, we inoculated GFP-K139 lysogens onto chitin particles to cultivate biofilm growth as described above. Biofilms were imaged once per day for 6 days. For each day, all cells were identified by constitutive expression of *mKate2*2, and cells that had induced to the lytic cycle were detected by the presence of GFP-labeled phage signal within *mKate2*2-positive cells. Lysogen induction frequency in co-culture was calculated as the 3D biovolume overlap between phage signal volume and lysogen signal volume, normalized to total lysogen biovolume (SI Appendix, Fig. S13). This metric was computed for lysogenic populations in monoculture and for lysogens in competition with naïve susceptible cells in co-culture.

#### Phage adsorption rate

Phage adsorption rate was calculated as previously described (97). Briefly, an overnight *Vibrio cholerae* culture was diluted and grown for 1 h at 37°C to 1-2×10^6^ cells/mL. Freshly prepared K139 phages (1×10^4^ PFU/ml) were added to aliquots of host bacteria, mixed by vortex, and incubated without shaking for 5 min at 37 °C. Two 1 mL samples were taken: one was centrifuged to separate adsorbed phage, and the second sample was left uncentrifuged. Supernatants were plated for plaque counts to determine un-adsorbed phage (N_0_) and total phage (N_t_) counts. Adsorption rate *d* was calculated as *N*_0_ = *N*_t_ *e*^-5*d*^

#### Image analysis

Zeiss CZI files were converted to .tiff stacks and imported into BiofilmQ v0.2.2 (65), a MATLAB-based custom image-processing framework for biofilm image data. Biovolume segmentation for bacterial cells and chitin was performed using semi-automated thresholding, with all segmentation cross-checked against the corresponding raw image data. For population dynamics experiments, bacterial biovolume was taken as a proxy for the population size of each bacterial strain. For experiments measuring lytic induction rate in the biofilm setting, segmented phage volume overlapping with segmented bacterial volume was taken as a proxy for the population size of lysogens undergoing lytic induction.

#### Replication and statistics

Mann-Whitney *U* tests were used for all pairwise comparisons. Biological replicates were defined as independent microfluidic device channels, with 1–3 chitin flakes per chamber serving as technical replicates. All plots were generated using Python with the Matplotlib library. Statistical analyses were performed using SciPy and statsmodels in Python. Python scripts were executed in Jupyter Notebook via Anaconda, with data handling facilitated by NumPy and Pandas.

#### Percolation model

To quantify how spatial structure and lysogen frequency influence the potential for phage-mediated killing, we implemented a spatial percolation framework in both two- and three-dimensional lattices. In the two-dimensional formulation, individual bacterial cells occupy sites on a square lattice; in the three-dimensional formulation, cells are arranged in a cubic lattice. Each site is assigned to one of three states: non-lysogen (phage-susceptible), lysogenic (immune), or infected. The simulation begins with the lattice fully occupied, with non-lysogen and lysogenic cells distributed randomly or in clustered distributions according to a predefined lysogen frequency. Cells remained fixed in place. At initialization, a single susceptible cell at the center of the lattice is designated as infected. Infection spreads deterministically: at each discrete time step, infected cells transmit to all adjacent side- and corner-facing neighbors (8 neighbors in 2D, 26 neighbors in 3D). A non-lysogen neighbor becomes infected, whereas lysogen blocks further transmission due to superinfection immunity. Lysogens do not change state and act as barriers to the spread of infection. The simulation proceeds until no further transmission events are possible. Simulations were implemented in Python using the NumPy and Matplotlib libraries for numerical computation and visualization, respectively, and the OS package was used for automated file handling. The model employed synchronous update rules and periodic boundary conditions. All model code is available for download on <u>GitHub</u>.

#### Population dynamics model

We supplemented the percolation modeling results with a spatial PDE reaction-diffusion model of phage interaction with susceptible non-lysogens and superinfection-immune lysogens (with parameter ranges supported by experiments where feasible, SI Appendix Fig. S12, Fig. S13). This model was used to assess the population dynamics of cells susceptible to infection (S), lysogenized and superinfection-immune cells (L), and free phage particles (P). The model builds on classical frameworks describing temperate phage dynamics (4–6, 98, 99) and incorporates both vertical transmission via lysogen growth and horizontal transmission via infection (4, 6, 99, 100). Susceptible cells and lysogens follow logistic growth to a shared carrying capacity *K*, each with equivalent maximum growth rate *r*.

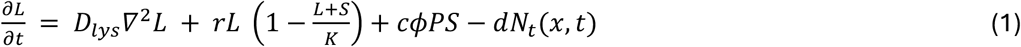

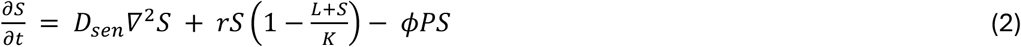

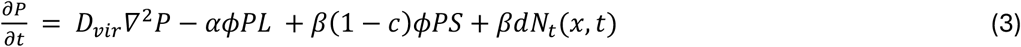

This model is a two-dimensional spatial reaction–diffusion simulation that tracks three populations on a grid. The spatial domain is represented as a (51 * 51) grid over a unit square, with each grid point containing local densities of (L), (S), and (P). Initial cell occupancy is randomly assigned with 10% of grid points occupied, and occupied sites are then assigned as either lysogens or susceptible cells according to the specified initial lysogen frequency. Phage-mediated interactions occur through adsorption, where *φ* denotes the adsorption rate of phages to host cells, including both susceptible cells and lysogens. After adsorption to a susceptible cell, a fraction *c* of infections results in genome integration and conversion of the susceptible cell into a lysogen, while the remaining infections follow the lytic pathway and produce new phages with burst size *β*. *N_t_*(*x*, *t*) denotes the number of lytic induction events at location *x* up to time *t*. Each induction event releases new phages, also with burst size *β*. Free phages can also be neutralized through adsorption to lysogens, which act as sinks for phage particles because of superinfection immunity. The parameter *α* scales the strength of this phage loss by representing the number of phage particles that can adsorb to superinfection-immune lysogens.

Simulations consisted of three steps for each iteration: diffusion; local interaction of lysogens, susceptible cells, and phages according to the reaction rules above; and finally the stochastic process of spontaneous lytic induction. Localized lytic induction is implemented as a Poisson process with *N_t_*(*x*, *t*) as the cumulative number of induction events at location x up to time t, a counting process with intensity *λ*(*x*, *t*) = *γL*(*x*, *t*); the increment *dN_t_*(*x*, *t*) is Poisson-distributed with mean *λ*(*x*, *t*)*dt*.

Simulations were implemented in MATLAB. Results shown in SI Fig. S9 were derived from simulations with the parameter values in the SI Appendix, Table S2. Spatial diffusion was solved on a two-dimensional grid using a Crank–Nicolson scheme with no-flux boundary conditions, while nonlinear local reaction terms were updated implicitly using a backward-Euler formulation solved by Newton iteration at each grid point. All code is available on GitHub.

#### Sensitivity analysis for adsorption rate and lytic induction

To evaluate how the selection regime for lysogens depends on phage adsorption and spontaneous lytic induction, we performed a two-parameter sweep using the spatial reaction–diffusion model. For every pair of lytic induction (*γ*) and adsorption rate (*φ*) parameters, we ran simulations using ten initial lysogen frequencies ranging from 0.01 to 0.99. Each simulation was initiated with 10% of grid sites occupied by lysogens or non-lysogens, and phage density was initialized at zero for correspondence with experimental conditions. To avoid assigning biological meaning to very small changes in population composition, lysogens were classified as having increased (decreased) in frequency if their relative abundance increased (decreased) by 10^-3^ relative to the starting condition. The selection regime for lysogens was designated as uniformly positive (negative) if they increased (decreased) in frequency from all starting conditions. A negative frequency-dependent selection regime was designated when the change in lysogen frequency shifted from positive values at lower initial lysogen frequencies to negative values at higher initial lysogen frequencies. Neutral competition between lysogens and non-lysogens was only observed for the case in which lytic induction rate was set to zero.

### SI Tables

**SI Table S1.** List of strains and reagents used in the study.

| Name / Strain# | Genotype | Reference |
| --- | --- | --- |
| **Vibrio cholerae** |  |  |
| TND3473 | K139<sup>+</sup>, $\Delta lacZ::Spec^R-lacI-P_{tac}$-mKO-κ, $\Delta VC1807::Cm^R-lacI-P_{tac}$-GFP | This study |
| TND3508 | $\Delta K139::Tm^R$, $\Delta lacZ::Spec^R-lacI-P_{tac}$-mKO-κ, $\Delta VC1807::Cm^R-lacI-P_{tac}$-GFP | This study |
| TND3433 | K139<sup>+</sup>, $\Delta lacZ::Spec^R-lacI-P_{tac}$-mKate22, $\Delta VC1807::Cm^R-lacI-P_{tac}$-GFP*::*parS*<sup>MT1</sup> | (101) |
| TND3348 | Orf18-msfGFPv2 (native locus), $\Delta VC1807::Kan^R$, $\Delta VCA0692::Tm^R$, $\Delta lacZ::Spec^R-P_{recN}$-GFP | This study |
| TND3741 | igVCA0265-VCA0266::*Carb*<sup>R</sup>-*P*<sub>orf17</sub>-Orf17-Orf18-msfGFPv2, $\Delta VCA0692::Tm^R$, $\Delta lacZ::Spec^R-lacI-P_{tac}$-mKate22 | This study |
| TND3742 | $\Delta K139::Tm^R$, $\Delta lacZ::Spec^R-lacI-P_{tac}$-mKO-κ | This study |
| CNV 304 | Orf18-msfGFPv2 (native locus), $\Delta VC1807::Kan^R$, $\Delta VCA0692::Tm^R$, $\Delta lacZ::Spec^R-lacI-P_{tac}$-mKate22 | This study |
| **E. coli K12** |  |  |
| CNE 748 | pBW-mKate22, *Kan*<sup>R</sup> | Lab stock |
| **Chemicals and Reagents** | **Source** | **Product number** |
| Poly-dimethylsiloxane (PDMS) | Dow Chemical Company SYLGARD 184 | cat. #04019862 |
| #1.5 glass coverslips | Azer Scientific | cat. #1152260 |
| Inlet tubing | Cole Palmer | cat. #06417-11 |
| 27Gx1/2 needles | BD Precision | cat. #30510 |
| 1mL syringes | Brandzig | cat. #CMD2583 |
| Harvard Apparatus Pico Plus Elite syringe pumps | Harvard Apparatus | cat. #70-4506 |
| Chitin flakes | Sigma-Aldrich | cat. #C9752 |
| N-Acetyl-D-glucosamine<br>(GlcNAc)<sub>1</sub> | Sigma-Aldrich | cat. #A8625 |
| Calcium Chloride | Sigma-Aldrich | cat. #C8106 |
| Magnesium Sulfate | Sigma-Aldrich | cat. #M2643 |
| Lysogeny Broth (LB), Miller | Sigma-Aldrich | cat. #L3552 |
| LB Agar, Miller | Sigma-Aldrich | cat. #L3147 |
| Sucrose | Sigma-Aldrich | cat. #S0389 |
| Yeast extract | Sigma-Aldrich | cat. #70161 |
| Agar powder | Thermo Scientific | cat. #A10752.36 |
| Sodium chloride | Fisher Scientific | cat. #S271 |
| Tryptone | Sigma-Aldrich | cat. #T7293 |
| Kanamycin | Sigma-Aldrich | cat. #K1377 |
| Polymyxin | Sigma-Aldrich | cat. #P0972 |
| **Software and Algorithms** | **Source** | **Version** |
| Zen Black | Zeiss | v14.0.0.0 |
| Zen Blue | Zeiss | v3.4.91.00000 |
| MATLAB | MathWorks | vR2023b |
| BiofilmQ | PMID: 33398098 (65) | v0.2.2 |
| Python | Python.org | v3.11.14 |
| Jupyter Notebook | Jupyter.org (102) | v6.5.4 |
| Anaconda | Anaconda.org(103) | V2.6.4 |
| NumPy | NumPy.org (104) | v2.3 |
| Pandas | Pandas.pydata.org(105) | v2.3.3 |
| Matplotlib | Matplotlib.org(106) | v3.10.7 |
| SciPy | Scipy.org(107) | v1.16.2 |
| statsmodels | Statsmodels.org (108) | v0.14.4 |

**SI Table S2.** Model parameters for SI Appendix Fig. S9.

| Variables |  |  | Description |
| --- | --- | --- | --- |
| L |  |  | normalized lysogen cell density |
| S |  |  | normalized susceptible cell density |
| P |  |  | normalized phage density |
| Parameters | Values | Unit | Description |
| $D_{lys}, D_{sen}$ | $10^{-7}$ | (domain length)<sup>2</sup> / simulation time | Diffusion coefficient for bacteria |
| $D_{vir}$ | $10^{-6}$ | (domain length)<sup>2</sup> / simulation time | Diffusion coefficient for free phages |
| $K$ | 1 | normalized density | Carrying capacity |
| $r$ | 0.1 | 1 / simulation time | Maximum growth rate cells. |
| $\phi$ | $[10^{-3}, 6 \times 10^{-1}]$ | 1 / (phage density * simulation time) | Adsorption of susceptible cells and phage |
| $c$ | 0.01 | dimensionless | Fraction of adsorption events that lead lysogenization of susceptible cells. |
| $\beta$ | 12 | phages / burst event | Number of phages added per lytic infection or lysogen induction event |
| $\alpha$ | 100 | dimensionless | Scaling factor for superinfection immunity |
| $\gamma$ | [0,200] | 1 / (lysogen density * simulation time) | Mean lytic induction rate per lysogen density |

### SI Figures

**SI Figure S1.**
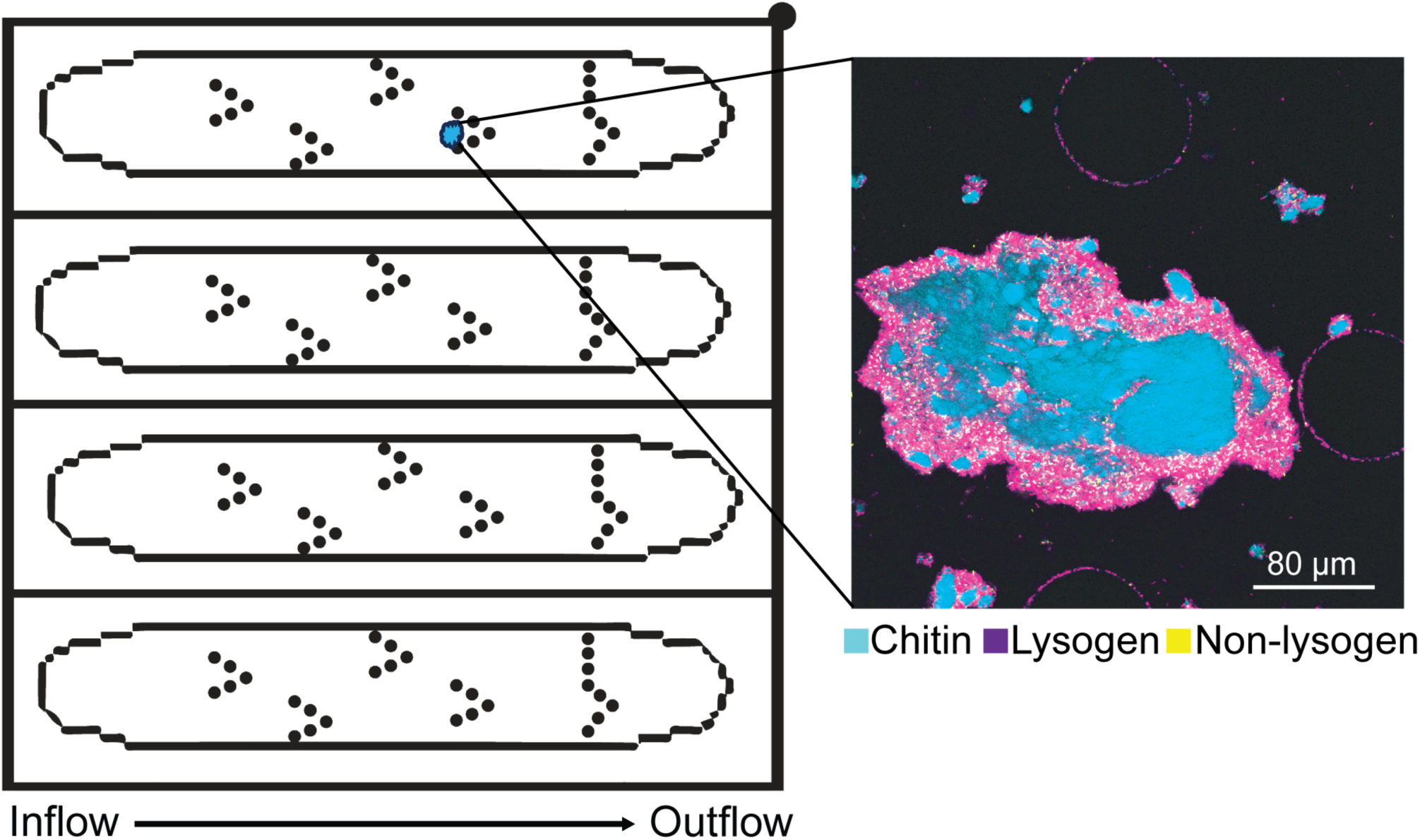
Schematic representation of the chitin-trapping microfluidic device. The diagram on the left illustrates a top-down view of the microfluidic chip design, with columns arranged in ’V’ orientations that trap chitin particles (cyan) in the flow path. The confocal image on the right shows a representative chitin particle (with bacteria inoculated) held in place by a column trap.

**SI Figure S2.**
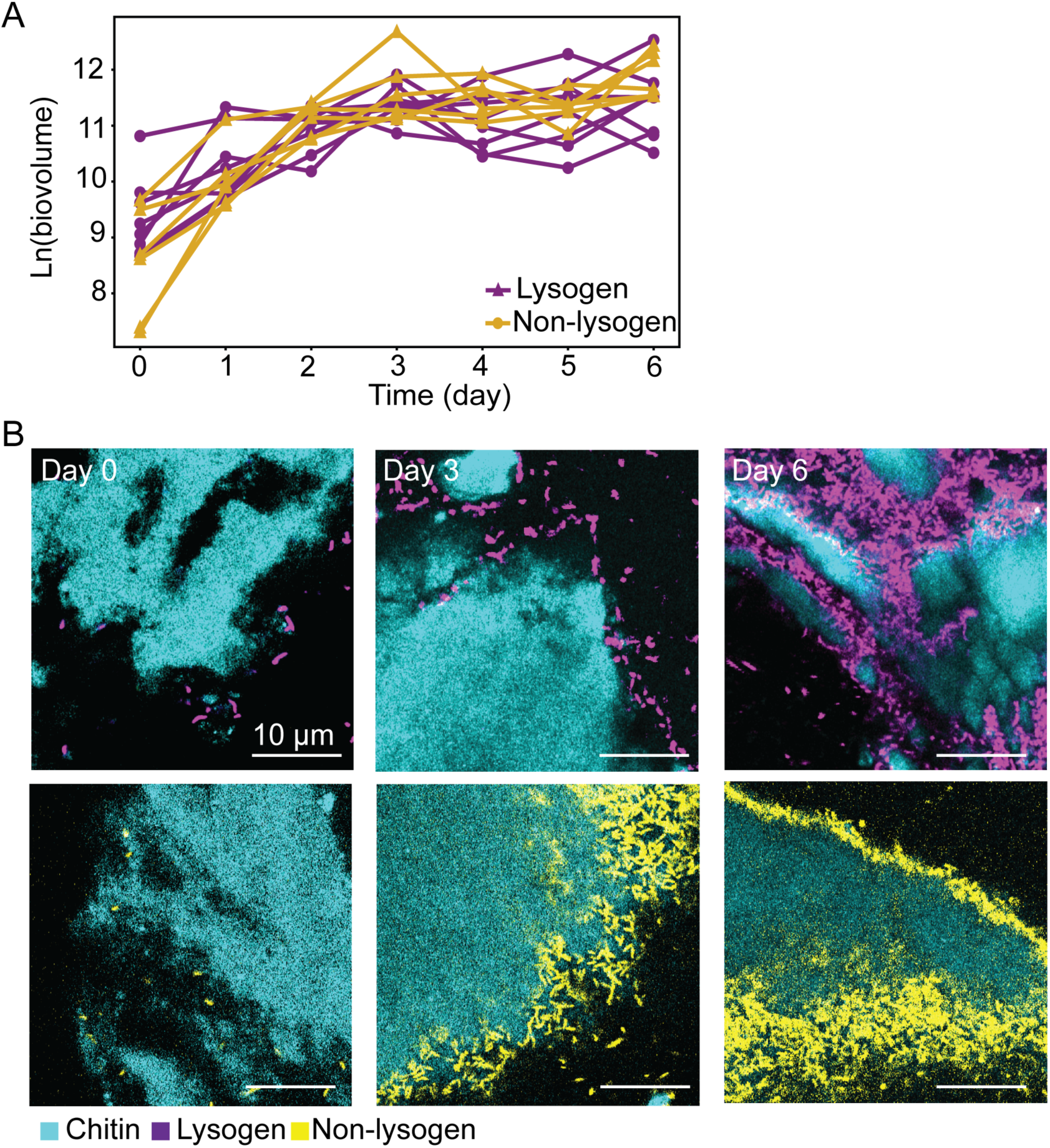
Growth and architectural dynamics of monoculture biofilm controls. (A) Time-series quantification of total biovolume for non-lysogen *V. cholerae* (*n* = 6) and K139-lysogenized *V. cholerae* (*n* = 7) monoculture biofilms over 7 days on chitin with continuous flow of seawater. Biovolume values are log-transformed. (B) Representative confocal images of biofilms from day 0, 3, and 5. Non-lysogen cells are shown in yellow; lysogens are shown in purple, and chitin is visible in blue.

**SI Figure S3.**
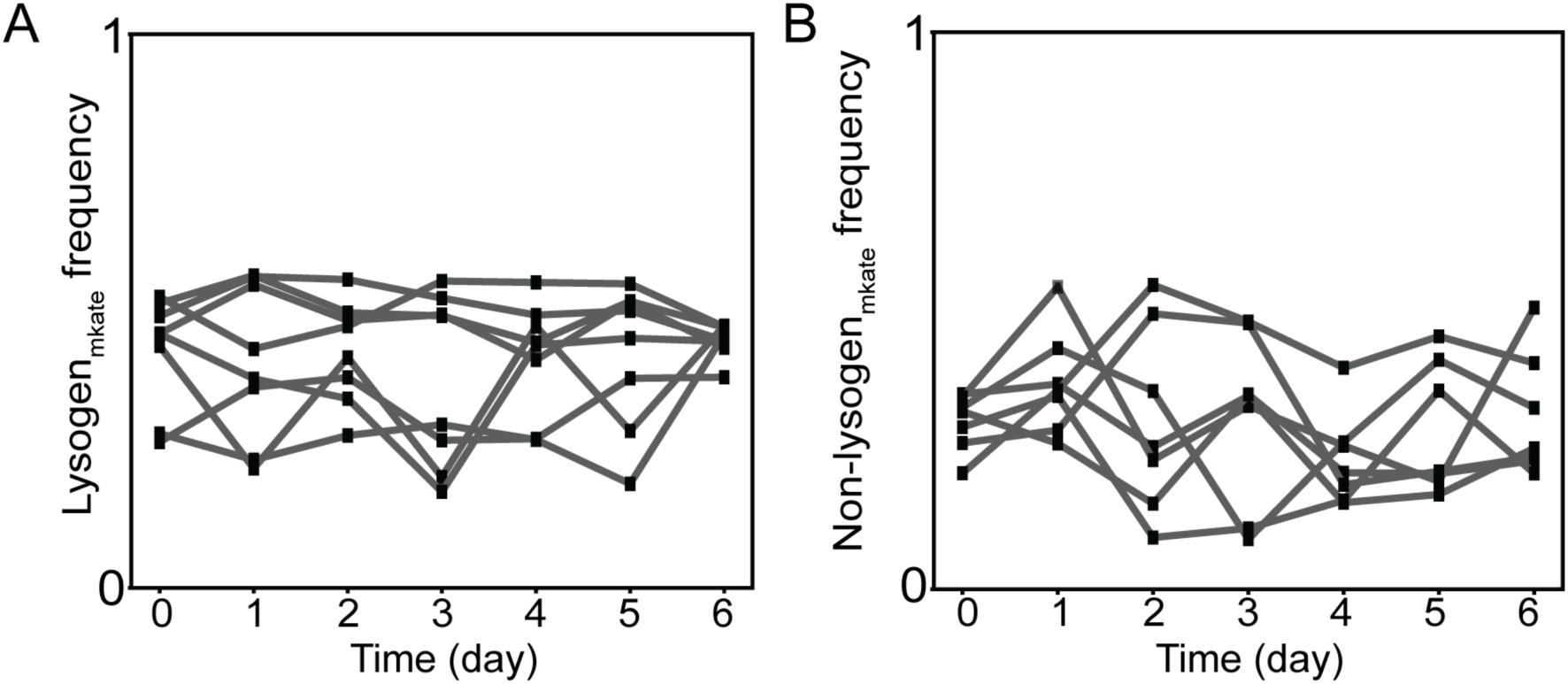
The different fluorescent protein variants are competitively neutral in biofilms on chitin. **(A)** Frequency of mKate2-labeled lysogens (left) and mKO-κ-labeled isogenic lysogens over a 6-day co-culture on chitin under flow. **(B)** Frequency of mKate2-labeled non-lysogens (left) and mKO-κ-labeled isogenic non-lysogens over a 6-day co-culture on chitin under flow.

**SI Figure S4.**
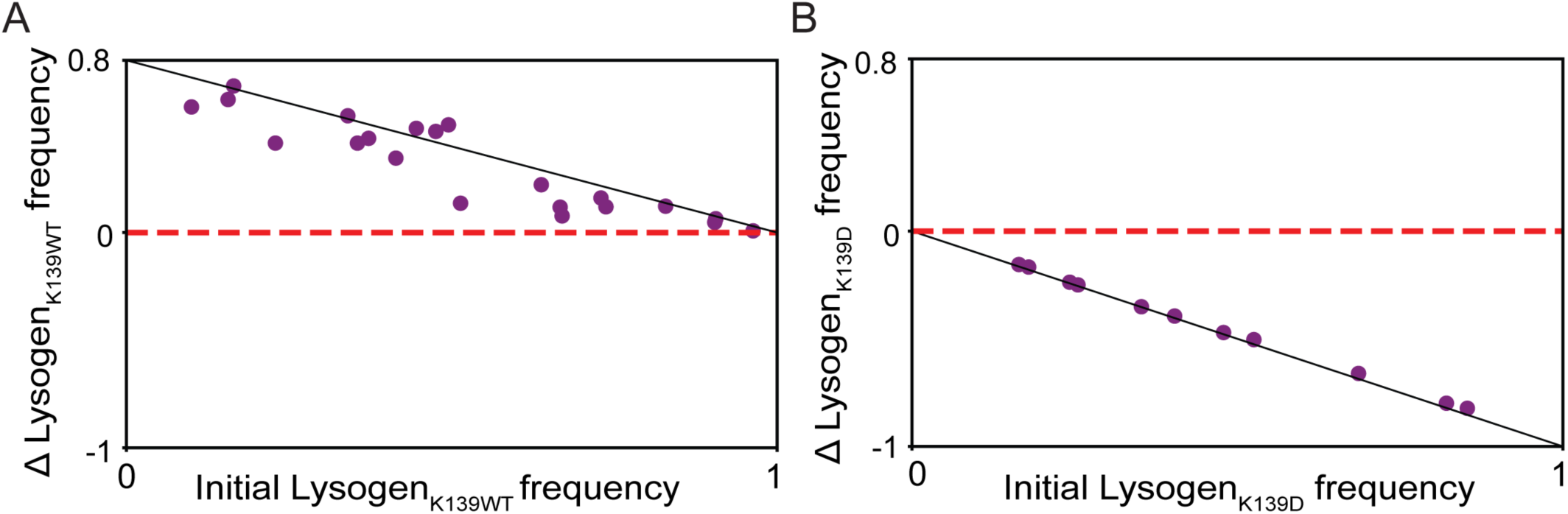
Lysogens that release viable phages are positively selected in liquid culture, but isogenic lysogens that release non-infectious phages are negatively selected. Lysogens harboring (A) wild type K139 prophages or (B) GFP-K139_D_ prophages were mixed with susceptible non-lysogens at a range of initial fractions in artificial seawater supplemented with 0.5% GlcNAc. Co-cultures were incubated with shaking at room temperature for 40 h. (A) When lysogens contained prophages that produce infectious virions after lytic induction, lysogens are positively selected from any initial condition. (B) When lysogens contained prophages that produce non-infectious phages after lytic induction, lysogens are negatively selected from any initial condition. Note that while the initial population density in liquid culture is similar to the density of host cells at the start of the main text biofilm experiments, the final population density in liquid culture was lower (local bacterial volume fraction: ∼0.1) than the final population density in the biofilm experiments (local bacterial volume fraction: ∼0.3).

**SI Figure S5.**
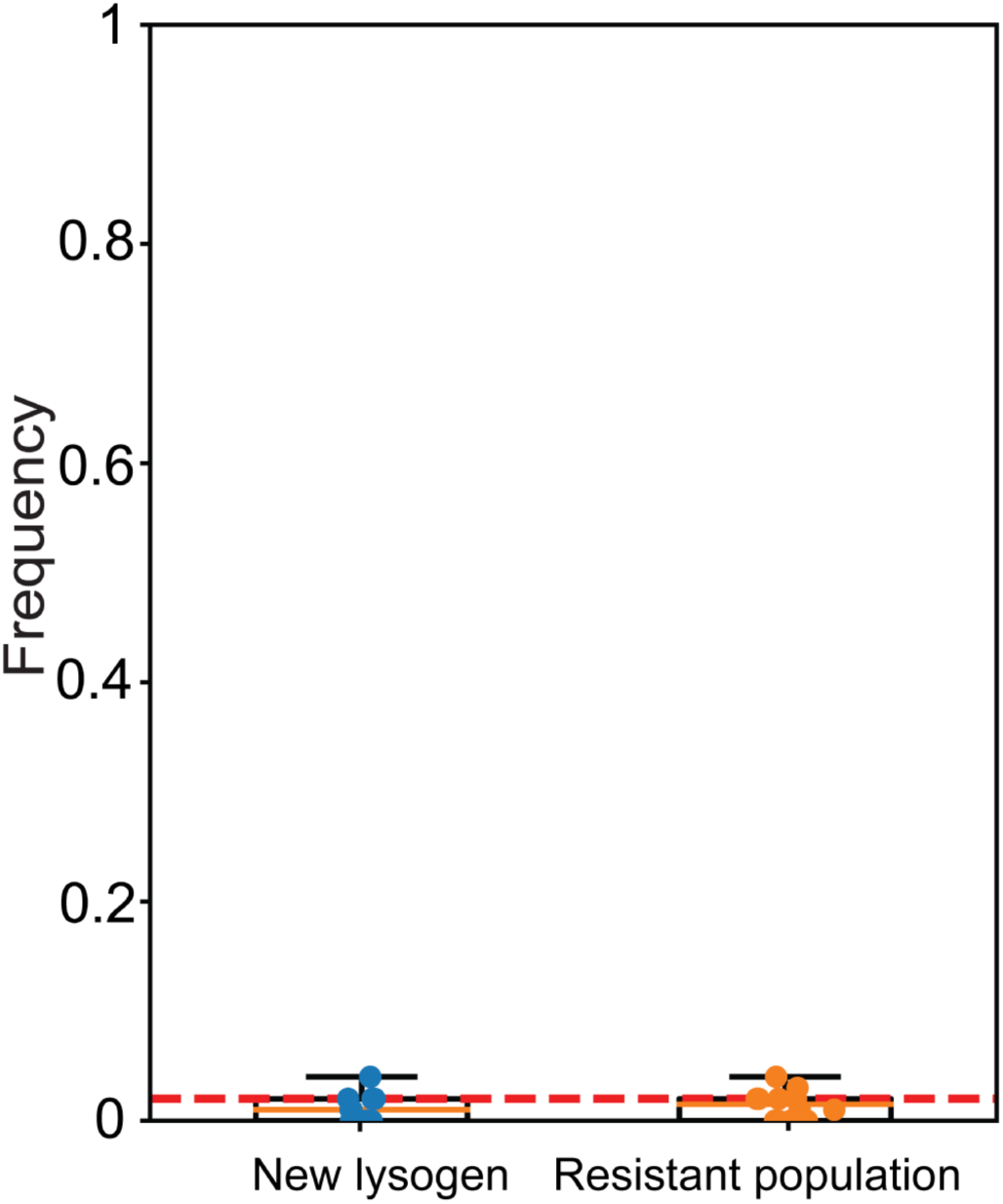
Novel lysogens and *de novo* phage-resistant mutants are rare in *V. cholerae* biofilms on chitin. Box plots depicting the frequency of newly lysogenized cells or *de novo* phage-resistant mutants among initially non-lysogenized, phage-susceptible *V. cholerae* after 6 days in co-culture with initially lysogenized *V. cholerae* on chitin. Initial lysogen frequencies for these experiments ranged 0.1 to 0.9 (*n* = 7). Each point represents the results of an independent chamber. The red line indicates the limit of detection.

**SI Figure S6.**
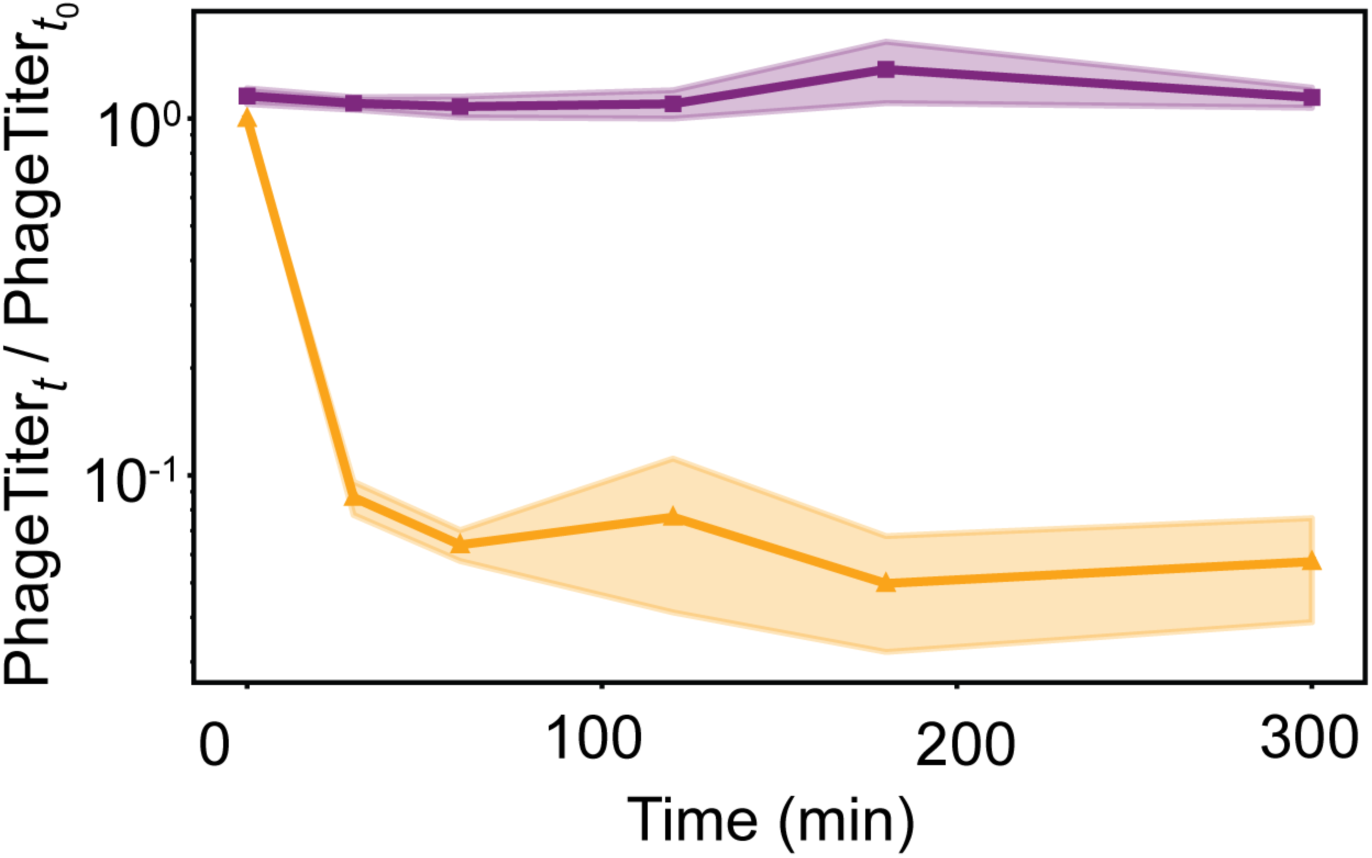
Phage adsorption assay in liquid culture with K139-lysogenized *V. cholerae* in comparison with a no-bacteria control. Phage titer (PFU/mL) over time (normalized to initial phage titer at *t*=0) following addition of purified K139 phages to cultures containing either K139-lysogenized *V. cholerae* at 1×10^5^ CFU/mL (shown in orange), or no bacteria (control, shown in purple). Cultures were incubated with continuous shaking in artificial seawater with 0.5% GlcNAc at 37° C. The shaded region denotes standard error of the mean (*n* = 3).

**SI Figure S7.**
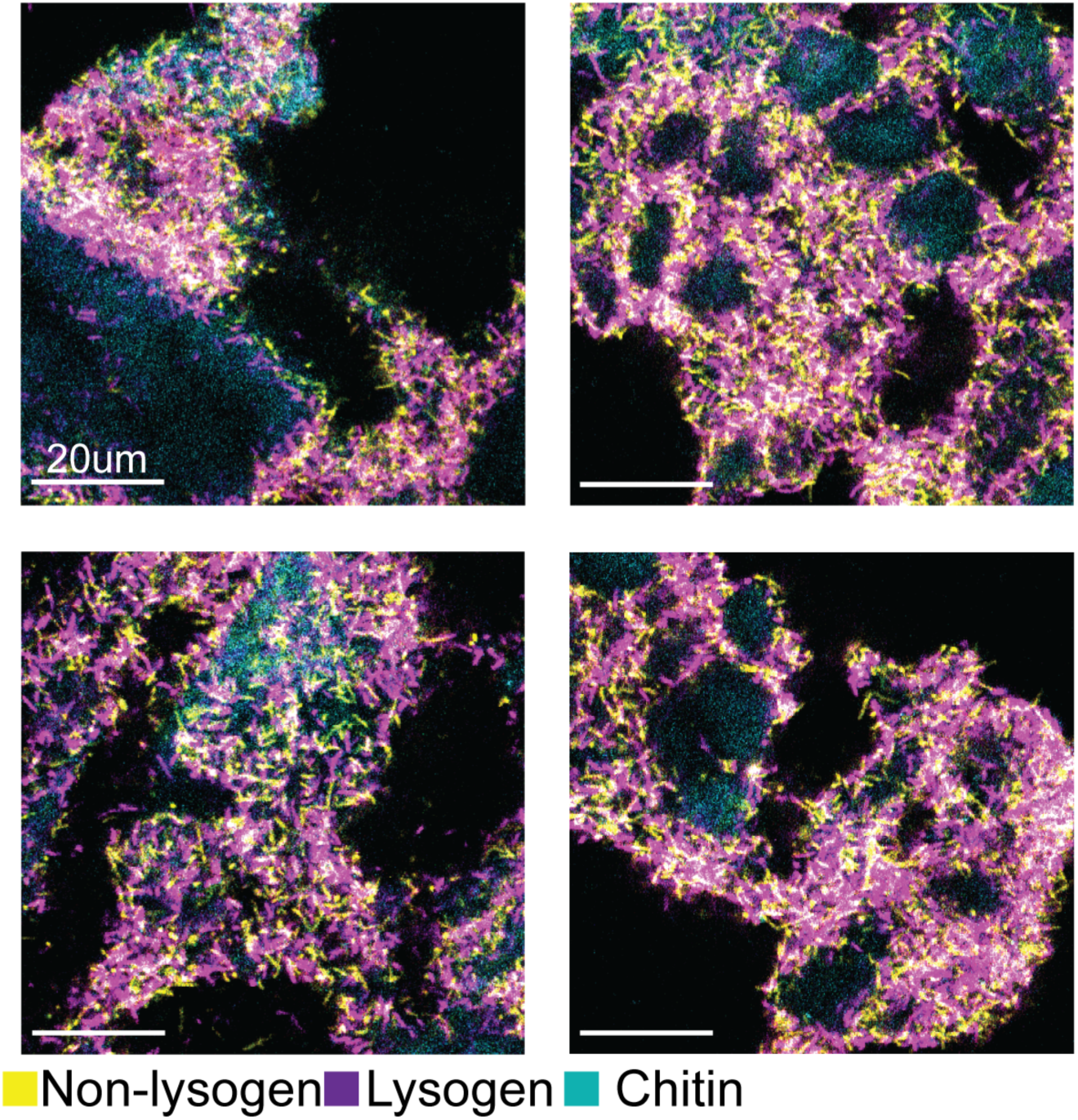
Lysogens and non-lysogens remain intermixed in biofilms on chitin surfaces. Four representative images of K139 lysogens (purple) and non-lysogens (yellow) after 6 days of biofilm growth on chitin. Lysogens were initialized at frequency 0.1-0.3. Note that these images are 2-dimensional optical sections through 3-dimensional chitin particles, with the two bacterial strains located along the particles water-exposed outer surface.

**SI Figure S8.**
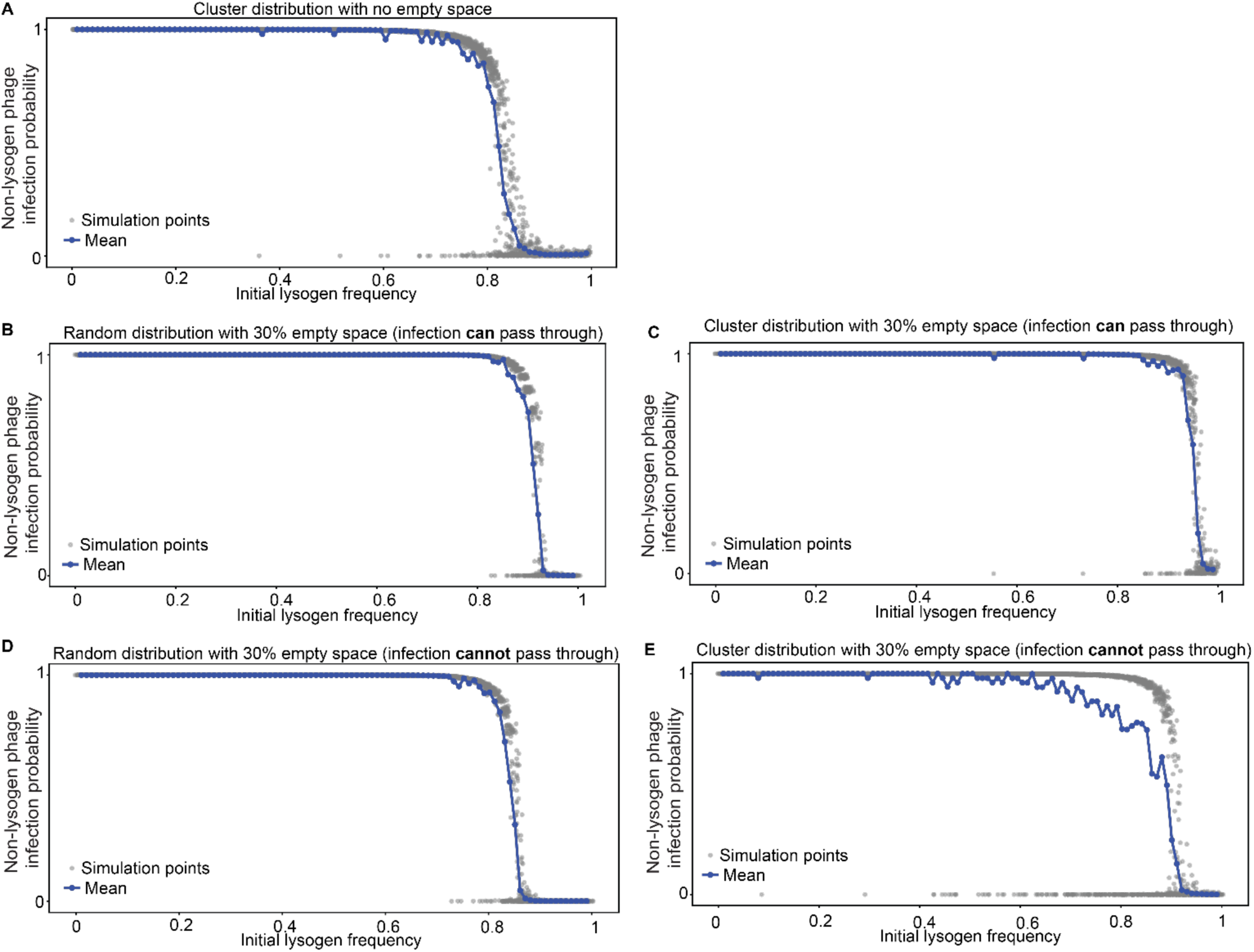
Alternative initial distributions of lysogens and non-lysogens yielded qualitatively similar thresholds of initial lysogen fraction at which the susceptible cells were no longer phage-exposed. The percolation model simulations from Figure 2 of the main text were repeated for several different scenarios of lysogen and non-lysogen arrangement to assess the robustness of model results to alternative initial conditions: (A) Clustered distributions of lysogens and non-lysogens; (B) Random distribution of lysogens and non-lysogens, including 30% open space through which phages infection is assumed to propagate freely. (C) Clustered distributions of lysogens and non-lysogens, including 30% open space through which phages infection is assumed to propagate freely. (D) Random distribution of lysogens and non-lysogens, including 30% open space through which phages infection cannot propagate. (E) Clustered distributions of lysogens and non-lysogens, including 30% open space through which phages infection cannot propagate.

**SI Figure S9.**
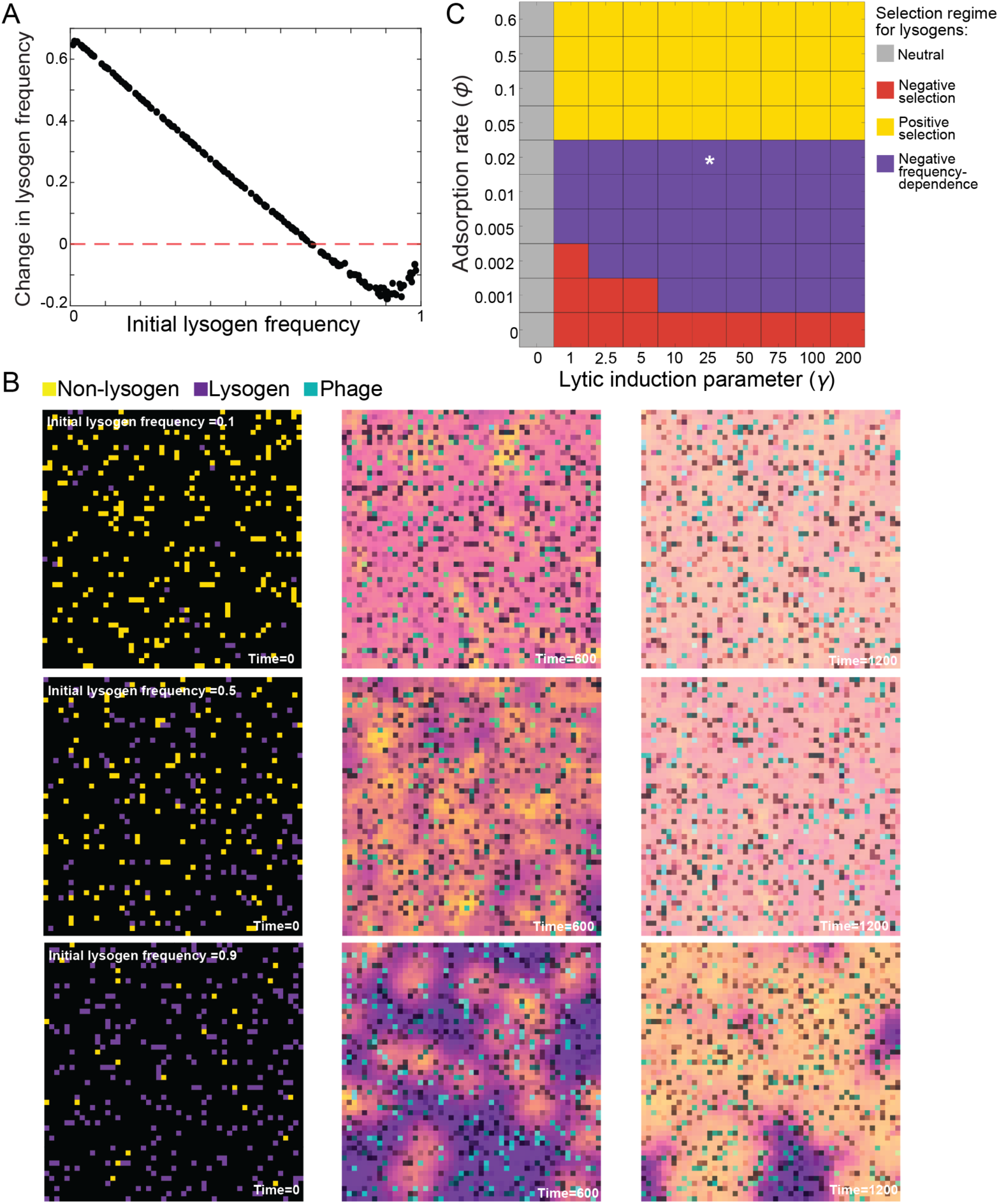
A PDE reaction-diffusion model recapitulates coexistence of lysogens and susceptible cells. (A) Example plot of the change in lysogen frequency as a function of initial lysogen frequency in the reaction-diffusion model of lysogen and non-lysogen population dynamics. (B) Representative frames from three simulations in panel A, corresponding to initial lysogen frequencies of 0.1, 0.5, and 0.9, from top to bottom row. The color channels are assigned directly from the code as cyan = free phage, purple = lysogens, and yellow = susceptible cells; overlapping populations can therefore appear as mixed colors depending on which signals coexist at the same grid position. (C) Selection regimes observed for different combinations of the adsorption rate and lytic induction parameter. For each grid square, simulations were run for a range of initial lysogen frequencies to determine if lysogens undergo uniformly positive selection, uniformly negative selection, negative frequency-dependent selection, or neutral competition with non-lysogens. The central grid square containing an asterisk corresponds to the parameter combination for example simulation results shown in panels A and B.

**SI Figure S10.**
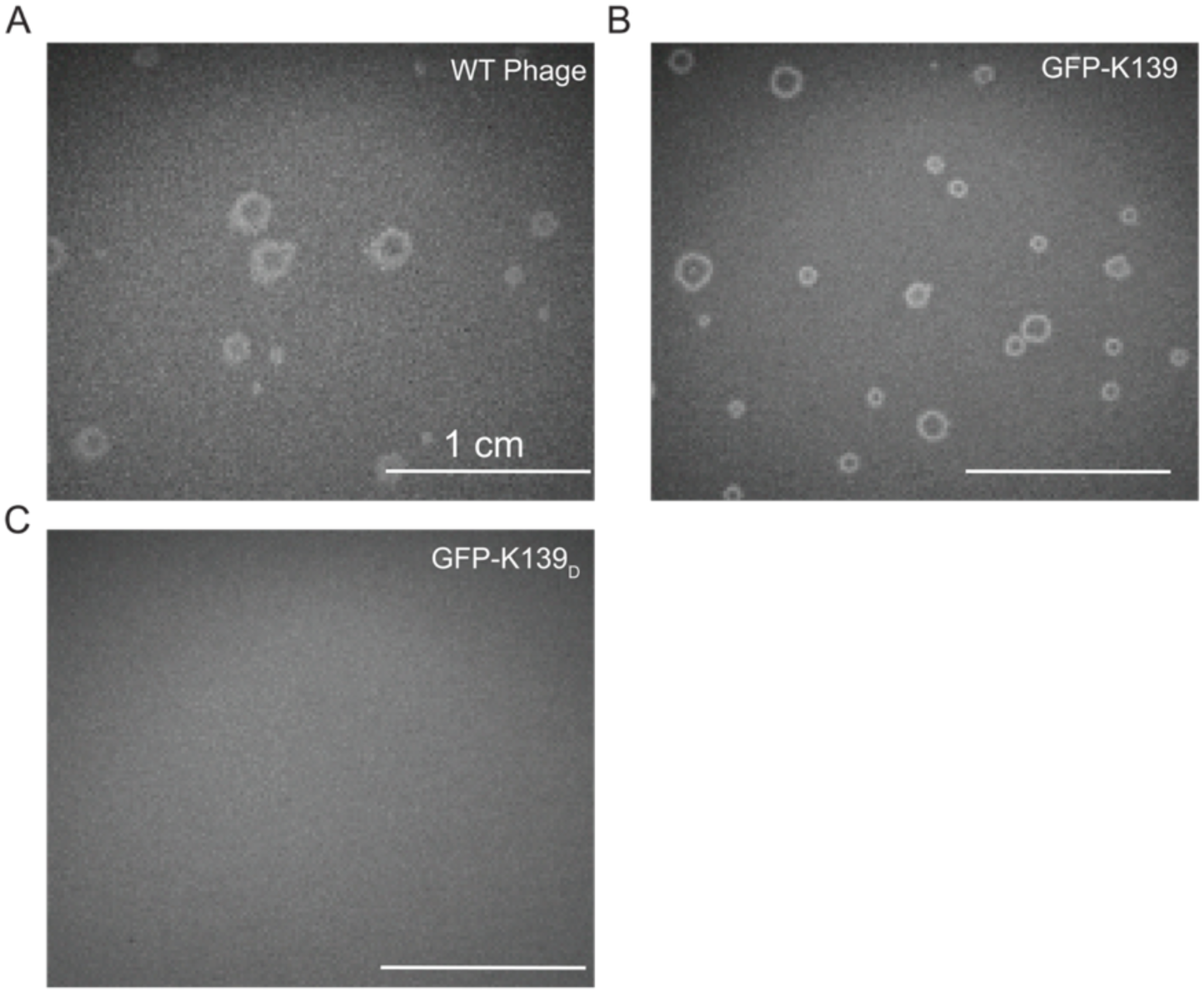
Plaque assays of wild-type, GFP labelled, and infection-defective phages on susceptible *Vibrio cholerae* lawns. Representative images of plaque formation following overnight incubation on lawns of phage-susceptible *V. cholerae* with phage lysates. **(A)** Wild-type K139 phage, showing clear zones of lysis. **(B)** Functional fluorescently labeled GFP-K139 phages, exhibiting comparable plaque morphology to wild-type. **(C)** Infection-defective GFP-K139_D_ phage, showing no detectable plaques.

**SI Figure S11.**
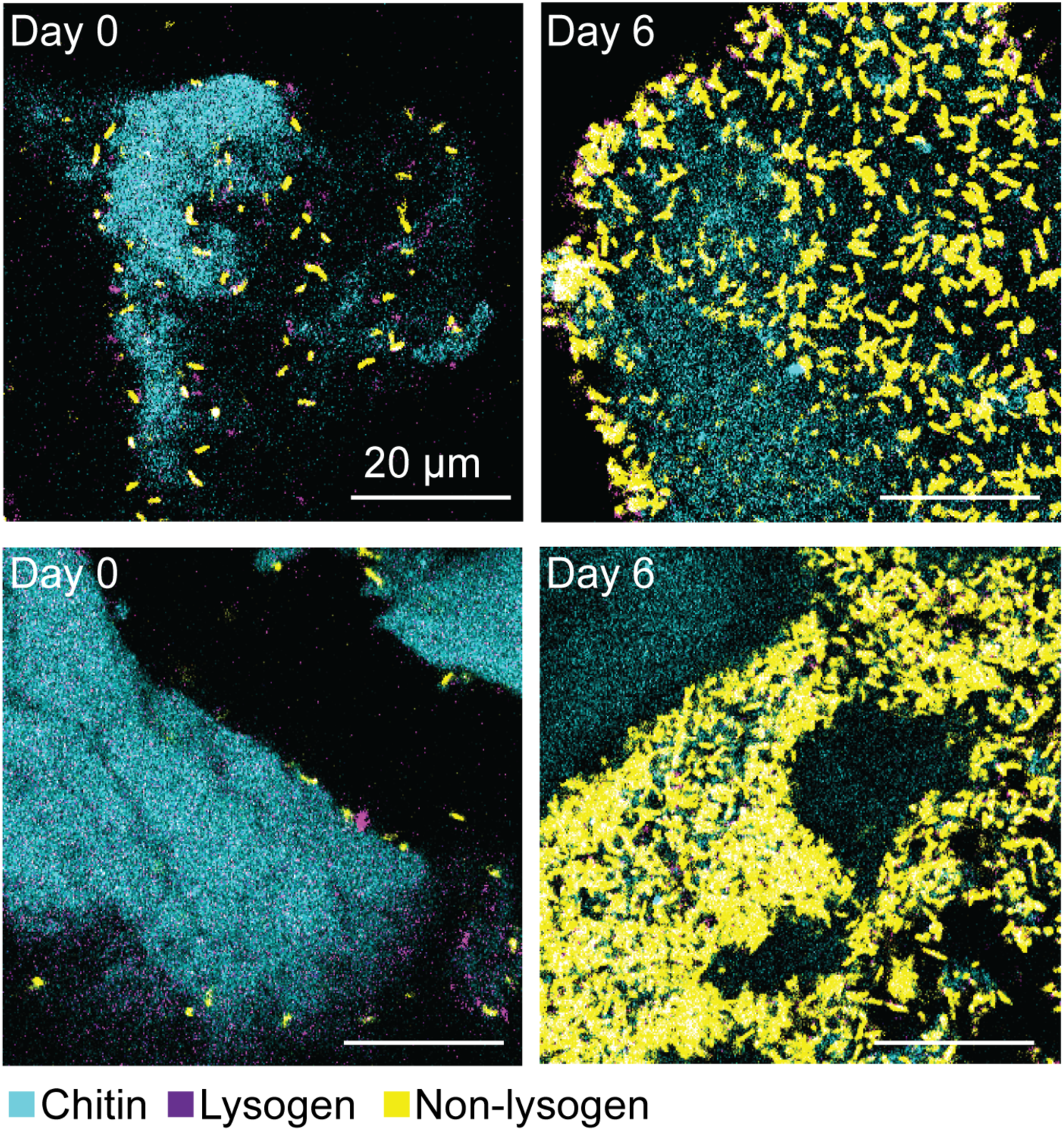
Representative images of lysogens carrying K139_D_ in co-culture with susceptible *V. cholerae* on chitin. Representative images at day 0 (D0) and day 6 (D6) showing susceptible cells (yellow), lysogens carrying non-functional K139_D_ phage (purple). In this condition, lysogens are progressively lost during competition, resulting in susceptible cells-dominated biofilms by day 6. These are representative images for quantitative data shown in Figure 3D in the Main Text.

**SI Figure S12.**
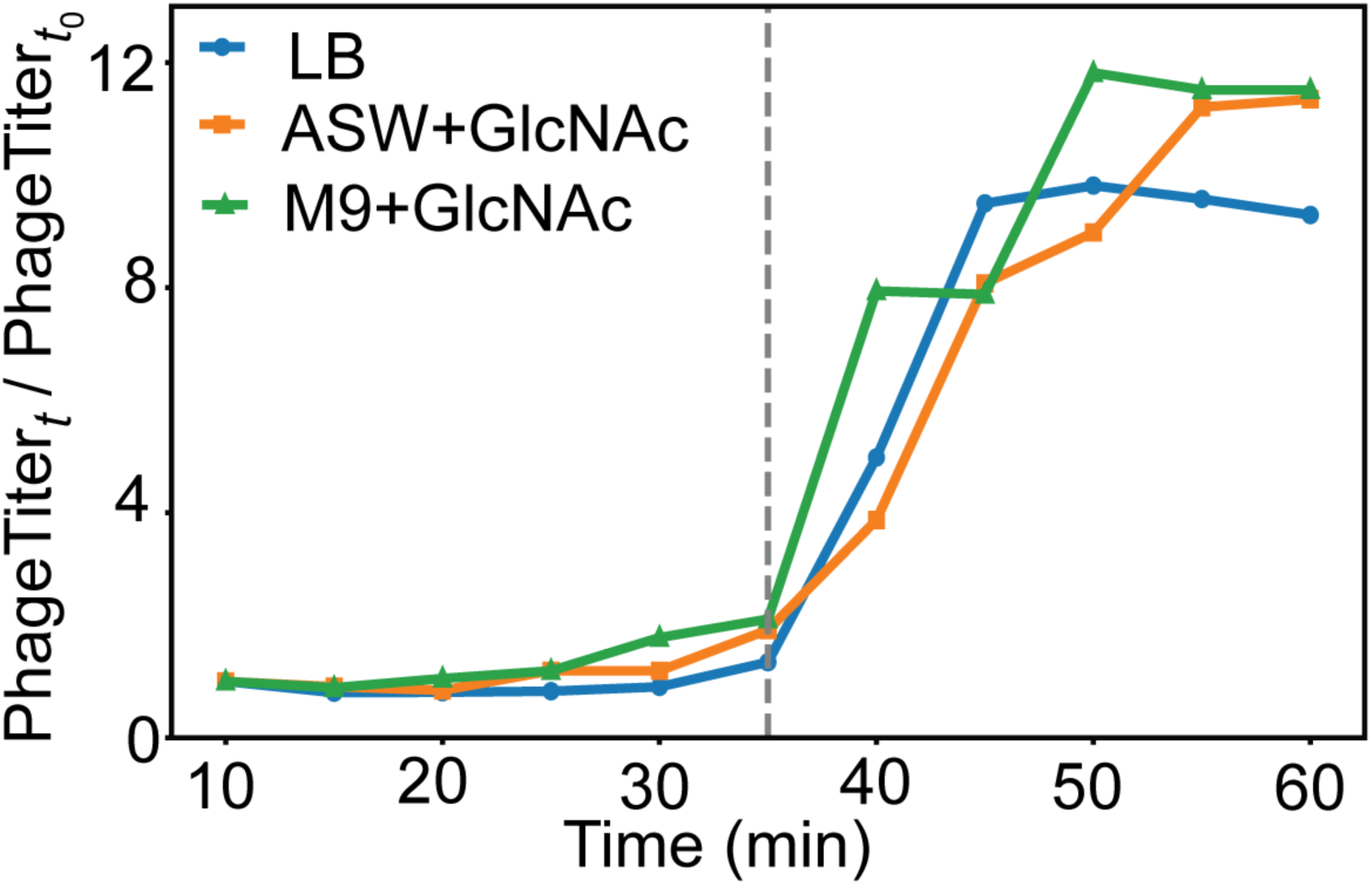
One-step growth curve of K139 phage in defined media. Infection cycles of K139 in co-culture with susceptible *V. cholerae* in LB medium, artificial seawater with 0.5% GlcNAc, and M9 minimal medium with 0.5% GlcNAc. Phage titer was measured via plaque assay every 5 minutes for 60 minutes. Estimated burst size was ∼12 PFU per cell; latency period ∼30–40 minutes. Data are averaged from two experiments per medium.

**SI Figure S13.**
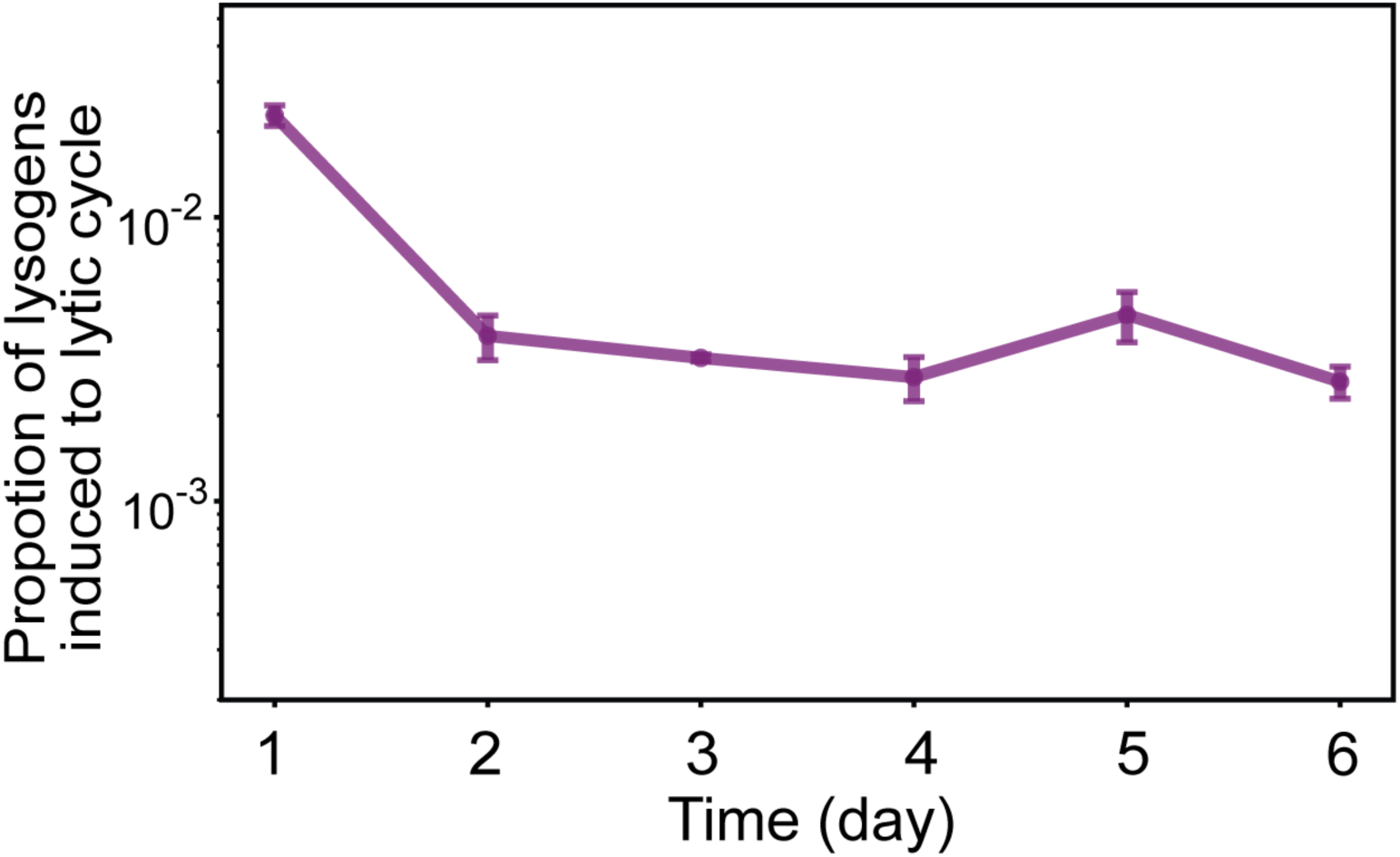
Proportion of lysogens undergoing phage induction in biofilms on chitin. Temporal trajectory of the proportion of lysogens undergoing lytic induction in biofilms grown on chitin flakes in flow (*n* = 3). Error bars represent standard errors of mean.

## References

1. Lwoff André, LYSOGENY. Bacteriol. Rev. 17, 269–337 (1953).

2. M. d’Herelle, Sur un microbe invisible antagoniste des bacilles dysentériques. Acta Kravsi (1961).

3. Barnhart B J, Cox S H, Jett J H, Prophage induction and inactivation by UV light. J. Virol. 18, 950–955 (1976).

4. F. M. Stewart, B. R. Levin, The population biology of bacterial viruses: Why be temperate. Theor. Popul. Biol. 26, 93–117 (1984).

5. J. S. Weitz, G. Li, H. Gulbudak, M. H. Cortez, R. J. Whitaker, Viral invasion fitness across a continuum from lysis to latency†. Virus Evol. 5, vez006 (2019).

6. G. Li, M. H. Cortez, J. Dushoff, J. S. Weitz, When to be temperate: on the fitness benefits of lysis vs. lysogeny. Virus Evol. 6, veaa042 (2020).

7. M. Ptashne, A genetic switch: phage λ and higher organisms. *No Title* (1992).

8. G. Ofir, R. Sorek, Contemporary Phage Biology: From Classic Models to New Insights. Cell 172, 1260–1270 (2018).

9. I. Herskowitz, D. Hagen, The lysis-lysogeny decision of phage λ: explicit programming and responsiveness. Annu. Rev. Genet. 14, 399–445 (1980).

10. B. Knowles, et al., Lytic to temperate switching of viral communities. Nature 531, 466–470 (2016).

11. M. De Paepe, et al., Carriage of λ Latent Virus Is Costly for Its Bacterial Host due to Frequent Reactivation in Monoxenic Mouse Intestine. PLOS Genet. 12, e1005861 (2016).

12. Nanda Arun M., Thormann Kai, Frunzke Julia, Impact of Spontaneous Prophage Induction on the Fitness of Bacterial Populations and Host-Microbe Interactions. J. Bacteriol. 197, 410–419 (2015).

13. C. Howard-Varona, K. R. Hargreaves, S. T. Abedon, M. B. Sullivan, Lysogeny in nature: mechanisms, impact and ecology of temperate phages. ISME J. 11, 1511–1520 (2017).

14. Bossi Lionello, Fuentes Juan A., Mora Guido, Figueroa-Bossi Nara, Prophage Contribution to Bacterial Population Dynamics. J. Bacteriol. 185, 6467–6471 (2003).

15. S. J. Labrie, J. E. Samson, S. Moineau, Bacteriophage resistance mechanisms. Nat. Rev. Microbiol. 8, 317–327 (2010).

16. Thomas M. J. N., Brockhurst M. A., Coyte K. Z., What makes a temperate phage an effective bacterial weapon? mSystems 9, e01036–23 (2024).

17. S. P. Brown, L. Le Chat, M. De Paepe, F. Taddei, Ecology of Microbial Invasions: Amplification Allows Virus Carriers to Invade More Rapidly When Rare. Curr. Biol. 16, 2048–2052 (2006).

18. X.-Y. Li, et al., Temperate phages as self-replicating weapons in bacterial competition. J. R. Soc. Interface 14, 20170563 (2017).

19. J. Haaber, et al., Bacterial viruses enable their host to acquire antibiotic resistance genes from neighbouring cells. Nat. Commun. 7, 13333 (2016).

20. E. Harrison, M. A. Brockhurst, Ecological and Evolutionary Benefits of Temperate Phage: What Does or Doesn’t Kill You Makes You Stronger. BioEssays 39, 1700112 (2017).

21. P. Hyman, S. T. Abedon, “Chapter 7 - Bacteriophage Host Range and Bacterial Resistance” in Advances in Applied Microbiology, (Academic Press, 2010), pp. 217–248.

22. K. D. Seed, et al., Phase Variable O Antigen Biosynthetic Genes Control Expression of the Major Protective Antigen and Bacteriophage Receptor in Vibrio cholerae O1. PLOS Pathog. 8, e1002917 (2012).

23. J. Bertozzi Silva, Z. Storms, D. Sauvageau, Host receptors for bacteriophage adsorption. FEMS Microbiol. Lett. 363, fnw002 (2016).

24. R. E. Lenski, Coevolution of bacteria and phage: Are there endless cycles of bacterial defenses and phage counterdefenses? J. Theor. Biol. 108, 319–325 (1984).

25. J. R. Meyer, et al., Repeatability and Contingency in the Evolution of a Key Innovation in Phage Lambda. Science 335, 428–432 (2012).

26. B. Koskella, M. A. Brockhurst, Bacteria–phage coevolution as a driver of ecological and evolutionary processes in microbial communities. FEMS Microbiol. Rev. 38, 916–931 (2014).

27. A. R. Hall, P. D. Scanlan, A. Buckling, Bacteria-Phage Coevolution and the Emergence of Generalist Pathogens. Am. Nat. 177, 44–53 (2011).

28. J. M. Borin, S. Avrani, J. E. Barrick, K. L. Petrie, J. R. Meyer, Coevolutionary phage training leads to greater bacterial suppression and delays the evolution of phage resistance. Proc. Natl. Acad. Sci. 118, e2104592118 (2021).

29. M. C. Bond, L. Vidakovic, P. K. Singh, K. Drescher, C. D. Nadell, Matrix-trapped viruses can prevent invasion of bacterial biofilms by colonizing cells. eLife 10, e65355 (2021).

30. E. V. Davies, et al., Temperate phages both mediate and drive adaptive evolution in pathogen biofilms. Proc. Natl. Acad. Sci. 113, 8266–8271 (2016).

31. S. T. Abedon, Bacteriophage exploitation of bacterial biofilms: phage preference for less mature targets? FEMS Microbiol. Lett. 363, fnv246 (2016).

32. C. D. Nadell, K. Drescher, N. S. Wingreen, B. L. Bassler, Extracellular matrix structure governs invasion resistance in bacterial biofilms. ISME J. 9, 1700–1709 (2015).

33. C. D. Nadell, J. B. Xavier, K. R. Foster, The sociobiology of biofilms. FEMS Microbiol. Rev. 33, 206–224 (2009).

34. J. W. Costerton, Z. Lewandowski, D. E. Caldwell, D. R. Korber, H. M. Lappin-Scott, MICROBIAL BIOFILMS. Annu. Rev. Microbiol. 49, 711–745 (1995).

35. H.-C. Flemming, et al., Biofilms: an emergent form of bacterial life. Nat. Rev. Microbiol. 14, 563–575 (2016).

36. E. L. Simmons, K. Drescher, C. D. Nadell, V. Bucci, Phage mobility is a core determinant of phage–bacteria coexistence in biofilms. ISME J. 12, 531–543 (2018).

37. C. D. Nadell, K. Drescher, K. R. Foster, Spatial structure, cooperation and competition in biofilms. Nat. Rev. Microbiol. 14, 589–600 (2016).

38. K. Drescher, C. D. Nadell, H. A. Stone, N. S. Wingreen, B. L. Bassler, Solutions to the Public Goods Dilemma in Bacterial Biofilms. Curr. Biol. 24, 50–55 (2014).

39. C. D. Nadell, et al., Cutting through the complexity of cell collectives. Proc. R. Soc. B Biol. Sci. 280, 20122770 (2013).

40. P. S. Stewart, M. J. Franklin, Physiological heterogeneity in biofilms. Nat. Rev. Microbiol. 6, 199–210 (2008).

41. Simmons Emilia L., et al., Biofilm Structure Promotes Coexistence of Phage-Resistant and Phage-Susceptible Bacteria. mSystems 5, 10.1128/msystems.00877-19 (2020).

42. W. W. Driscoll, J. W. Pepper, Theory for the evolution of diffusible external goods. Evolution 64, 2682–2687 (2010).

43. J. B. Winans, L. Zeng, C. D. Nadell, Spatial propagation of temperate phages within and among biofilms. Proc. Natl. Acad. Sci. 122, e2417058122 (2025).

44. J. B. Winans, B. R. Wucher, C. D. Nadell, Multispecies biofilm architecture determines bacterial exposure to phages. PLOS Biol. 20, e3001913 (2022).

45. A. Penesyan, I. T. Paulsen, S. Kjelleberg, M. R. Gillings, Three faces of biofilms: a microbial lifestyle, a nascent multicellular organism, and an incubator for diversity. Npj Biofilms Microbiomes 7, 80 (2021).

46. H. Sentenac, A. Loyau, J. Leflaive, D. S. Schmeller, The significance of biofilms to human, animal, plant and ecosystem health. Funct. Ecol. 36, 294–313 (2022).

47. K. Sauer, et al., The biofilm life cycle: expanding the conceptual model of biofilm formation. Nat. Rev. Microbiol. 20, 608–620 (2022).

48. B. A. Berryhill, T. Gil-Gil, A. P. Smith, B. R. Levin, The future of phage therapy in the USA. Trends Mol. Med. 31, 982–991 (2025).

49. K. Moon, et al., Considerations and perspectives on phage therapy from the transatlantic taskforce on antimicrobial resistance. Nat. Commun. 16, 10883 (2025).

50. J.-P. Pirnay, et al., Personalized bacteriophage therapy outcomes for 100 consecutive cases: a multicentre, multinational, retrospective observational study. Nat. Microbiol. 9, 1434–1453 (2024).

51. L. Vidakovic, P. K. Singh, R. Hartmann, C. D. Nadell, K. Drescher, Dynamic biofilm architecture confers individual and collective mechanisms of viral protection. Nat. Microbiol. 3, 26–31 (2018).

52. M. F. Hansen, S. L. Svenningsen, H. L. Røder, M. Middelboe, M. Burmølle, Big Impact of the Tiny: Bacteriophage–Bacteria Interactions in Biofilms. Trends Microbiol. 27, 739–752 (2019).

53. Conner Jenna G., Teschler Jennifer K., Jones Christopher J., Yildiz Fitnat H., Staying Alive: Vibrio cholerae’s Cycle of Environmental Survival, Transmission, and Dissemination. Microbiol. Spectr. 4, 10.1128/microbiolspec.vmbf-0015-2015 (2016).

54. M. Blokesch, Chitin colonization, chitin degradation and chitin-induced natural competence of Vibrio cholerae are subject to catabolite repression. Environ. Microbiol. 14, 1898–1912 (2012).

55. K. L. Meibom, et al., The Vibrio cholerae chitin utilization program. Proc. Natl. Acad. Sci. 101, 2524–2529 (2004).

56. C. Pruzzo, L. Vezzulli, R. R. Colwell, Global impact of Vibrio cholerae interactions with chitin. Environ. Microbiol. 10, 1400–1410 (2008).

57. Huq A, et al., Ecological relationships between Vibrio cholerae and planktonic crustacean copepods. Appl. Environ. Microbiol. 45, 275–283 (1983).

58. C. A. Hayes, T. N. Dalia, A. B. Dalia, Systematic genetic dissection of chitin degradation and uptake in Vibrio cholerae. Environ. Microbiol. 19, 4154–4163 (2017).

59. M. M. Omand, R. Govindarajan, J. He, A. Mahadevan, Sinking flux of particulate organic matter in the oceans: Sensitivity to particle characteristics. Sci. Rep. 10, 5582 (2020).

60. A. L. Alldredge, M. W. Silver, Characteristics, dynamics and significance of marine snow. Prog. Oceanogr. 20, 41–82 (1988).

61. Grossart Hans-Peter, Kiørboe Thomas, Tang Kam, Ploug Helle, Bacterial Colonization of Particles: Growth and Interactions. Appl. Environ. Microbiol. 69, 3500–3509 (2003).

62. M. Trucksis, J. Michalski, Y. K. Deng, J. B. Kaper, The Vibrio cholerae genome contains two unique circular chromosomes. Proc. Natl. Acad. Sci. 95, 14464–14469 (1998).

63. Kapfhammer Dagmar, Blass Julia, Evers Stefan, Reidl Joachim, Vibrio cholerae Phage K139: Complete Genome Sequence and Comparative Genomics of Related Phages. J. Bacteriol. 184, 6592–6601 (2002).

64. Nesper Jutta, Blaß Julia, Fountoulakis Michael, Reidl Joachim, Characterization of the Major Control Region ofVibrio cholerae Bacteriophage K139: Immunity, Exclusion, and Integration. J. Bacteriol. 181, 2902–2913 (1999).

65. R. Hartmann, et al., Quantitative image analysis of microbial communities with BiofilmQ. Nat. Microbiol. 6, 151–156 (2021).

66. G. L. Harrow, et al., Negative frequency-dependent selection and asymmetrical transformation stabilise multi-strain bacterial population structures. ISME J. 15, 1523–1538 (2021).

67. B. R. Levin, et al., Frequency-dependent selection in bacterial populations. Philos. Trans. R. Soc. Lond. B Biol. Sci. 319, 459–472 (1997).

68. C. Moore, M. E. J. Newman, Epidemics and percolation in small-world networks. *Phys*. Rev. E 61, 5678–5682 (2000).

69. M. Á. Serrano, M. Boguñá, Percolation and Epidemic Thresholds in Clustered Networks. Phys Rev Lett 97, 088701 (2006).

70. E. Kenah, J. C. Miller, Epidemic Percolation Networks, Epidemic Outcomes, and Interventions. Interdiscip. Perspect. Infect. Dis. 2011, 543520 (2011).

71. P. Grassberger, On the critical behavior of the general epidemic process and dynamical percolation. Math. Biosci. 63, 157–172 (1983).

72. H.-C. Flemming, et al., The biofilm matrix: multitasking in a shared space. Nat. Rev. Microbiol. 21, 70–86 (2023).

73. K. W. Bayles, Bacterial programmed cell death: making sense of a paradox. Nat. Rev. Microbiol. 12, 63–69 (2014).

74. G. R. Squyres, D. K. Newman, Single-cell lysis patterns morphogenesis of eDNA in the matrix of Pseudomonas aeruginosa biofilms. Proc. Natl. Acad. Sci. 122, e2514210122 (2025).

75. C. M. Carmody, J. M. Goddard, S. R. Nugen, Bacteriophage Capsid Modification by Genetic and Chemical Methods. Bioconjug. Chem. 32, 466–481 (2021).

76. E. Pavoni, P. Vaccaro, V. D’Alessio, R. De Santis, O. Minenkova, Simultaneous display of two large proteins on the head and tail of bacteriophage lambda. BMC Biotechnol. 13, 79 (2013).

77. V. L. Taylor, et al., Prophages block cell surface receptors to preserve their viral progeny. Nature 644, 1049–1057 (2025).

78. T. Britton, F. Ball, P. Trapman, A mathematical model reveals the influence of population heterogeneity on herd immunity to SARS-CoV-2. Science 369, 846–849 (2020).

79. T. Hiraoka, Z. Ghadiri, A. K. Rizi, M. Kivelä, J. Saramäki, Strength and weakness of disease-induced herd immunity in networks. Proc. Natl. Acad. Sci. 122, e2421460122 (2025).

80. F. Ball, L. Critcher, P. Neal, D. Sirl, The impact of household structure on disease-induced herd immunity. J. Math. Biol. 87, 83 (2023).

81. P. Payne, L. Geyrhofer, N. H. Barton, J. P. Bollback, CRISPR-based herd immunity can limit phage epidemics in bacterial populations. eLife 7, e32035 (2018).

82. S. Heilmann, K. Sneppen, S. Krishna, Coexistence of phage and bacteria on the boundary of self-organized refuges. Proc. Natl. Acad. Sci. 109, 12828–12833 (2012).

83. O. Kimchi, Y. Meir, N. S. Wingreen, Lytic and temperate phage naturally coexist in a dynamic population model. ISME J. wrae093 (2024). 10.1093/ismejo/wrae093.

84. B. Walton, et al., A biofilm-tropic Pseudomonas aeruginosa bacteriophage uses the exopolysaccharide Psl as receptor. eLife 13, RP102352 (2025).

85. Maffei Enea, Manner Christina, Jenal Urs, Harms Alexander, Complete genome sequence of Pseudomonas aeruginosa phage Knedl. Microbiol. Resour. Announc. 13, e01174–23 (2024).

86. L. E. Knecht, M. Veljkovic, L. Fieseler, Diversity and Function of Phage Encoded Depolymerases. Front. Microbiol. **Volume** 10**-**2019 (2020).

87. A. Visnapuu, M. Van der Gucht, J. Wagemans, R. Lavigne, Deconstructing the Phage–Bacterial Biofilm Interaction as a Basis to Establish New Antibiofilm Strategies. Viruses 14 (2022).

88. B. Rosan, R. J. Lamont, Dental plaque formation. Microbes Infect. 2, 1599–1607 (2000).

89. S. Wang, et al., Microbial Richness of Marine Biofilms Revealed by Sequencing Full-Length 16S rRNA Genes. Genes 13 (2022).

90. W. Zhang, et al., Marine biofilms constitute a bank of hidden microbial diversity and functional potential. Nat. Commun. 10, 517 (2019).

91. M. Castledine, A. Buckling, Critically evaluating the relative importance of phage in shaping microbial community composition. Trends Microbiol. 32, 957–969 (2024).

92. Q. Li, et al., Formation of Multispecies Biofilms and Their Resistance to Disinfectants in Food Processing Environments: A Review. J. Food Prot. 84, 2071–2083 (2021).

93. A. B. Dalia, E. McDonough, A. Camilli, Multiplex genome editing by natural transformation. Proc. Natl. Acad. Sci. 111, 8937–8942 (2014).

94. A. M. Kropinski, A. Mazzocco, T. E. Waddell, E. Lingohr, R. P. Johnson, “Enumeration of Bacteriophages by Double Agar Overlay Plaque Assay” in Bacteriophages: Methods and Protocols, Volume 1: Isolation, Characterization, and Interactions, M. R. J. Clokie, A. M. Kropinski, Eds. (Humana Press, 2009), pp. 69–76.

95. E. L. Ellis, M. Delbrück, The grorth of bacteriophage. J. Gen. Physiol. 22, 365–384 (1939).

96. A. M. Kropinski, “Practical Advice on the One-Step Growth Curve” in Bacteriophages: Methods and Protocols, Volume 3, M. R. J. Clokie, A. M. Kropinski, R. Lavigne, Eds. (Springer New York, 2018), pp. 41–47.

97. R. H. Heineman, J. J. Bull, Testing optimality with experimental evolution: lysis time in a bacteriophage. Evolution 61, 1695–1709 (2007).

98. T. W. Berngruber, R. Froissart, M. Choisy, S. Gandon, Evolution of Virulence in Emerging Epidemics. PLOS Pathog. 9, e1003209 (2013).

99. B. R. Levin, F. M. Stewart, L. Chao, Resource-Limited Growth, Competition, and Predation: A Model and Experimental Studies with Bacteria and Bacteriophage. Am. Nat. 111, 3–24 (1977).

100. Z. Qiu, The analysis and regulation for the dynamics of a temperate bacteriophage model. Math. Biosci. 209, 417–450 (2007).

101. J. D. Holt, Y. Peng, T. N. Dalia, A. B. Dalia, C. D. Nadell, Environmental DNA adsorption to chitin can promote horizontal gene transfer by natural transformation. Proc. Natl. Acad. Sci. 122, e2420708122 (2025).

102. T. Kluyver, et al., Jupyter Notebooks – a publishing format for reproducible computational workflows in Positioning and Power in Academic Publishing: Players, Agents and Agendas, F. Loizides, B. Schmidt, Eds. (IOS Press, 2016), pp. 87–90.

103. Anaconda Software Distribution. Anaconda Doc. (2020).

104. S. van der Walt, S. C. Colbert, G. Varoquaux, The NumPy Array: A Structure for Efficient Numerical Computation. Comput. Sci. Eng. 13, 22–30 (2011).

105. W. McKinney, others, Data structures for statistical computing in python in Proceedings of the 9th Python in Science Conference, (Austin, TX, 2010), pp. 51–56.

106. J. D. Hunter, Matplotlib: A 2D Graphics Environment. Comput. Sci. Eng. 9, 90–95 (2007).

107. P. Virtanen, et al., SciPy 1.0: fundamental algorithms for scientific computing in Python. Nat. Methods 17, 261–272 (2020).

108. S. Seabold, J. Perktold, statsmodels: Econometric and statistical modeling with python in *9th Python in Science Conference*, (2010).

